# Dense Computer Replica of Cortical Microcircuits Unravels Cellular Underpinnings of Auditory Surprise Response

**DOI:** 10.1101/2020.05.31.126466

**Authors:** Oren Amsalem, James King, Michael Reimann, Srikanth Ramaswamy, Eilif Muller, Henry Markram, Israel Nelken, Idan Segev

**Affiliations:** Department of Neurobiology, The Hebrew University of Jerusalem, Jerusalem, Israel; Blue Brain Project, École Polytechnique Fédérale de Lausanne, Geneva, Switzerland; The Edmond and Lily Safra Center for Brain Sciences, The Hebrew University of Jerusalem, Jerusalem, Israel

**Keywords:** Compartmental modeling, cortical signaling, stimulus-specific-adaptation, synaptic depression, spike frequency adaptation, emergence cortical dynamics

## Abstract

The nervous system is notorious for its strong response evoked by a surprising sensory input, but the biophysical and anatomical underpinnings of this phenomenon are only partially understood. Here we utilized *in-silico* experiments of a biologically-detailed model of a neocortical microcircuit to study stimulus specific adaptation (SSA) in the auditory cortex, whereby the neuronal response adapts significantly for a repeated (“expected”) tone but not for a rare (“surprise”) tone. SSA experiments were mimicked by stimulating tonotopically-mapped thalamo-cortical afferents projecting to the microcircuit; the activity of these afferents was modeled based on our *in-vivo* recordings from individual thalamic neurons. The modeled microcircuit expressed naturally many experimentally-observed properties of SSA, suggesting that SSA is a general property of neocortical microcircuits. By systematically modulating circuit parameters, we found that key features of SSA depended on synergistic effects of synaptic depression, spike frequency adaptation and recurrent network connectivity. The relative contribution of each of these mechanisms in shaping SSA was explored, additional SSA-related experimental results were explained and new experiments for further studying SSA were suggested.

## Introduction

Neurons in the primary auditory cortex, A1, exhibit a phenomenon called stimulus specific adaptation, SSA. When animals are presented with a tone sequence composed of a frequent tone (“standard”) and an occasional rare tone (“deviant”) the response of A1 neurons is markedly reduced to the standard tone, whereas their response to the same tone when deviant remains strong^1–5^ (see Nelken, 2014 for a definition and introduction to auditory SSA).

SSA is believed to emerge *de novo* in A1 as it is weak in the main thalamic input station to A1, the ventral medial geniculate body (MGBv)^6,9^. Therefore, A1 has become a prime focus for a number of experimental and theoretical studies of SSA^5,10–13^. Modeling studies suggest that SSA can emerge from feed-forward synaptic depression in the thalamocortical synapses^13–16^ and from local neocortical synaptic depression^11,17,18^. The models differ in their biological realism, ranging from population-based firing-rate models to spiking point-neuron-based models. Models which incorporate recurrent synaptic depression^11^ have the greatest success in replicating various properties of SSA, including the dependence of the magnitude of SSA on the frequency difference between the deviant and the standard, on the probability of the deviant, on the intertone interval, and on the input amplitude. These models also replicated the important property of specific deviance sensitivity – the larger response to a deviant tone (when the other tone presentations have a single expected tone frequency) than to the same tone, with the same probability, when the other tones vary in frequency. However, none of these models included the detailed cellular and synaptic diversity of neocortical microcircuits, nor did they replicate or explain the full repertoire of SSA without requiring specific parameter tuning. Indeed, we still lack a complete understanding of the underlying biophysical mechanisms of SSA. Towards this end, the present study uses a biologically-detailed dense computer reconstruction of a neocortical microcircuit, NMC^19^, to study SSA. This model, which has been extensively validated against anatomical and physiological experiments^20–23^ was challenged with the question: Does SSA as found in A1 emerge in this microcircuit without parameter adjustment? If it does, then such a model might have strong explanatory power for the underlying biophysical mechanisms of the phenomenon.

The NMC comprises a ∼0.3 mm^3^ volume of the neocortical tissue and is modeled after the rat somatosensory cortex, consisting of 6 layers, a total of ∼31,000 neurons, 55 morphological cell types, as well as the electrical properties of these neurons and of their synaptic connections (∼36 million synapses). The circuit was reconstructed based on the available experimental data and connectivity rules as described in ^19^ and ^24^. The model was not built in order to answer any specific *a priori* question, but to recreate the detailed anatomy and physiology of the NMC. Nevertheless, the model has been successfully used to perform *in silico* experiments that explore the anatomy and physiology of local neocortical circuits. For example, the model revealed the strong constraints of neuron morphology on local connectivity^25^ and how they result in non-random structures^26^, characterized the role of directed cliques of neurons in shaping neuronal responses^27^, and suggested mechanisms to suppress response variability caused by synaptic noise^28^.

We found that, when supplying the NMC with a biologically plausible auditory input via tonotopically-arranged thalamo-cortical axons, the model exhibited SSA with properties that mimicked many experimental observations. By modifying a variety of circuit parameters, we identified multiple mechanisms that may support SSA. In agreement with previous studies, synaptic depression of thalamo-cortical synapses as well as, to a lesser degree, depression of cortico-cortical synapses, shape SSA. Importantly, the model uncovered two additional features, spike frequency adaptation (SFA) and network connectivity that, even in the absence of synaptic depression, synergistically give rise to key features of SSA. This “replicate and explain” paradigm enabled us to explore the relative contribution of each of these mechanisms in determining the strength of SSA in the 6 neocortical layers and to replicated and explain other properties of SSA - its dependence on tone intensity^29,30^ and on neuron frequency preference^31^.

## Results

### Response to auditory surprise emerges naturally in the modeled thalamo-cortical circuit

The simulated dense neocortical microcircuit, NMC, consists of 31,346 neurons and ∼36 million intrinsic synapses (**Fig. 1a**). This microcircuit is innervated by 574 thalamo-cortical axons (**Fig. 1b** and see **Methods**), making a total of ∼ 2 million thalamo-cortical synapses. In this model, neurons in lower layer 3 and lower layer 5 receive the largest number of thalamo-cortical synapses^19^ in accordance with experimental data^32^. Each thalamo-cortical axon branches in the 3D volume of the NMC, creating multiple excitatory synapses on nearby cortical cells. An example of the postsynaptic cells receiving excitatory synaptic inputs from two thalamo-cortical axons (red and blue axons and respective colored cortical neurons) is shown in **Fig. 1c**. The hexagon at the bottom illustrates the tonotopic organization of the simulated thalamic input (see also hexagon at the middle of **Fig. 1d**).

**Figure 1.**
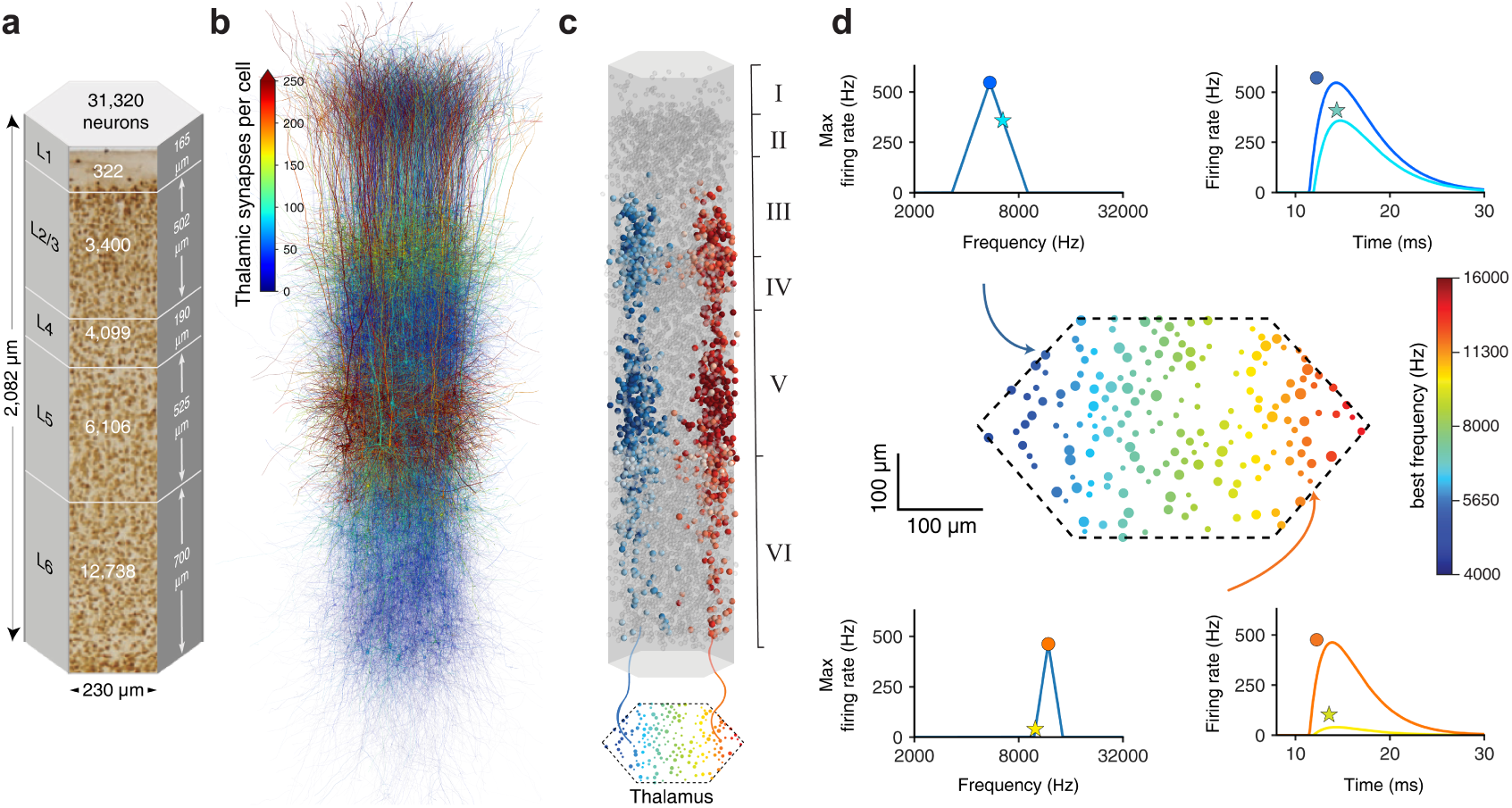
Feeding auditory-like input to the reconstructed cortical microcircuit. **a**. Dimensions and cell numbers per layer in the modeled microcircuit. The total number of neurons in this circuit is shown at the top. **b.** Number of thalamic synapses per cortical neuron; colored scale bar at top left. **c**. Tonotopic organization of the thalamic afferents is shown by colored dots in the lower hexagon. Two exemplar thalamo-cortical axons are shown, blue afferent with best frequency of 5,447 Hz and red afferent with best frequency of 11,854 Hz. The spatial distribution of the post-synaptic cortical neurons receiving synapses from these two axons is shown by the respective blue and red circles. **d**. Middle hexagon depicts the tonotopic organization of the thalamic inputs, color-coded by their best frequency. The size of each circle represents the width of the afferent’s tuning curve. A total of 574 thalamic afferents were simulated (only some are shown; see **Methods**). Top left shows the tuning curve for the afferent marked by the blue arrow. Top right shows the response of that afferent to two different tones (the best frequency, BF, marked by circle and for a 0.2 octave higher marked by an star). Bottom left and right, as in top traces for the thalamic afferent marked by the orange arrow. The modeled responses of the thalamic axons were all fitted to the experimental results shown in **Supplementary Fig. 1**.

To mimic auditory input impinging on the NMC from the thalamus, the time-dependent firing rate of the modeled thalamic axons was fitted to the rates measured experimentally in the auditory thalamus, MGBv (**Supplementary Fig. 1** and see **Methods**). For each simulated tone presented, the instantaneous firing rate, *f*(*t, s*), of each axon followed an alpha function Eq. (1). *f*(*t, s*) depend on the tuning curve of the axon FR(*s, b*), which has a triangular waveform Eq. (2). In those functions, *t* is time, *b* is the axon’s best frequency, and *s* is the tone frequency (see **Methods**). The parameters for Eq. (1,2) were fitted such that the simulated response of the thalamic axons to specific tone frequencies mimicked the experimentally measured response at tone intensity of 60 dB (**Methods** and **Supplementary Fig. 1**). Two exemplar tuning curves are shown in **Fig. 1d**, top and bottom frames at left, with their corresponding instantaneous firing rate shown respectively at right (*f*(*t, s*) for 2 input frequency; *s*). The modeled thalamic axons were organized in a tonotopic manner such that their best frequency spans 2 octaves over 560 μm (**Fig. 1d** hexagon in the middle, see details in **Methods**), in accordance with experimental findings^33,34^. Axons with low best frequency are marked by a bluish color and axons with high best frequency with a reddish color (color code scale at right).

Using the model illustrated in **Figure 1**, we next simulated the oddball experimental paradigm^1,13^ as follows. We selected two frequencies, *f*_*1*_ = 6,666 Hz and *f*_*2*_ = 9,600 Hz, that are centered (on log scale) around 8,000 Hz and presented them to the NMC with a 300 ms inter-stimulus interval. In one case, *f*_*1*_ served as a standard tone (presented 95% of the time) interspersed randomly 5% of the time with *f*_*2*_, the deviant tone. In the second case, the roles of *f*_*1*_ and *f*_*2*_ were reversed (see **Methods** and^13^). We focused our analysis on excitatory neurons located within ± 25 μm from the center of the simulated microcircuit (“center microcolumn”, see green neurons in **Fig. 4a**). These cortical neurons receive most of their thalamic synapses from axons whose best frequency is ∼8,000 Hz (green axons in the hexagon at **Fig. 1d**).

**Figure 2.**
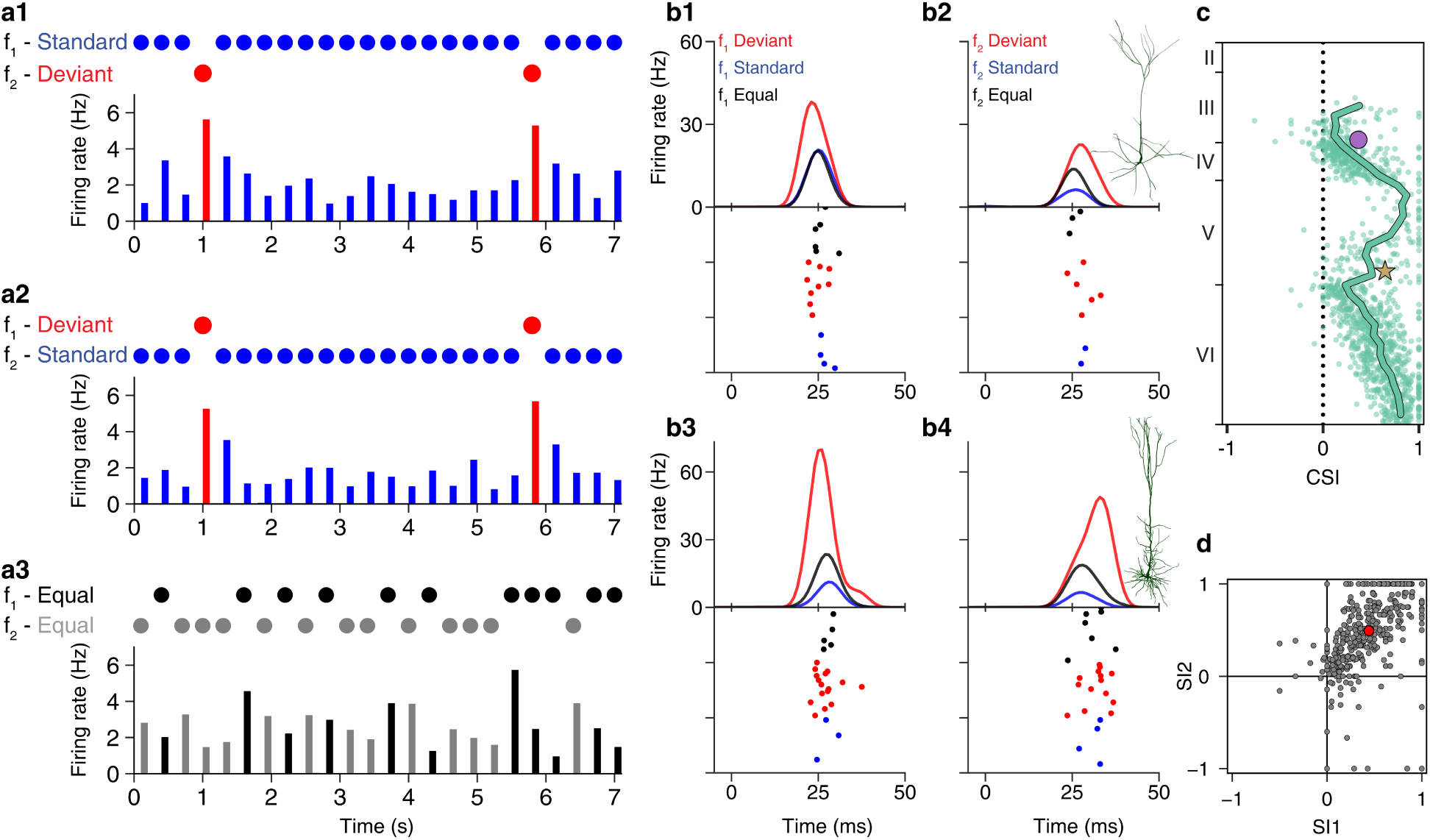
Stimulus-specific adaptation emerges in the modeled neocortical microcircuit. **a1-a3**. Blue and red bars - average response to the SSA protocol of all (2,696) responsive excitatory neurons located within ± 25 µm from the center of the simulated circuit (see **Methods**). **a1**. The standard frequency was *f*_*1*_ = 6,666 Hz (95% of the tones presented, blue circles) and the deviant frequency *f*_*1*_ = 9,600 Hz (red circles). **a2.** *f*_*2*_ was the standard and *f*_*1*_ the deviant. **a3.** *f*_*1*_ and *f*_*2*_ were presented with an equal probability. In all cases, the bin size was 100 ms. **b1-b4**. Response of two selected pyramidal cells shown in the inset (B1-B2, L23 PC; B3-B4, L5 TTPC1). In all cases, the response of the neurons to the deviant tone (red PSTH trace at tope and red raster below) was larger than when the same tone was the standard tone (blue PSTH and raster) or when *f*_*1*_ and *f*_*2*_ were represented equally (black PSTH and raster). The rasters of the standard and equal conditions show 25 randomly selected tone presentations (out of 475 standard presentations and 250 equal presentations). Each PSTH trace was smoothed by a 10 ms Hamming window. **c**. CSI of 1,350 excitatory neurons (green circles) and its mean across layers (thick green line). Circle and star denote the two selected cells shown in b. **d**. SI1 against SI2 for the neurons in c; both indices are mostly positive, implying that the majority of neurons respond more strongly to the deviant. The mean is marked by the red circle.

**Figure 3.**
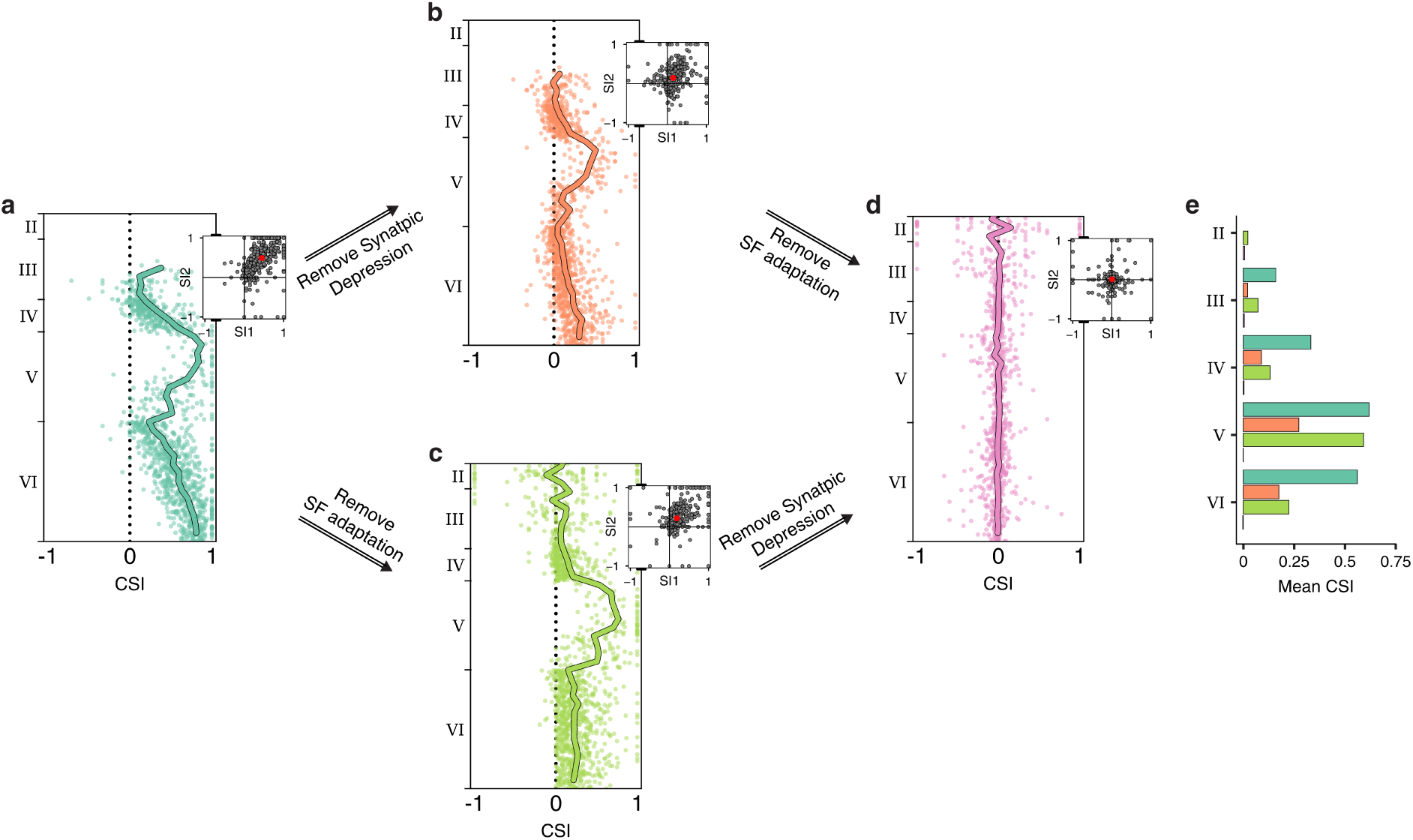
The contribution of synaptic depression and spike frequency adaptation to SSA. **a**. CSI and SI1 vs. SI2 (inset) indices for the excitatory neurons in the microcircuit as in **Fig. 2c. b**. As in a but removing synaptic depression in both thalamo-cortical and cortico-cortical synapses. The SSA is reduced but not abolished. **c**. Removing spike frequency adaptation (SFA) in all modeled neurons. Albeit reduced, the SSA persists. **d**. Removal of both SFA and synaptic depression eliminated the SSA. **e**. Mean CSI/layer for each of the microcircuit configurations shown in a-d. Colors are as in a-d.

**Figure 4.**
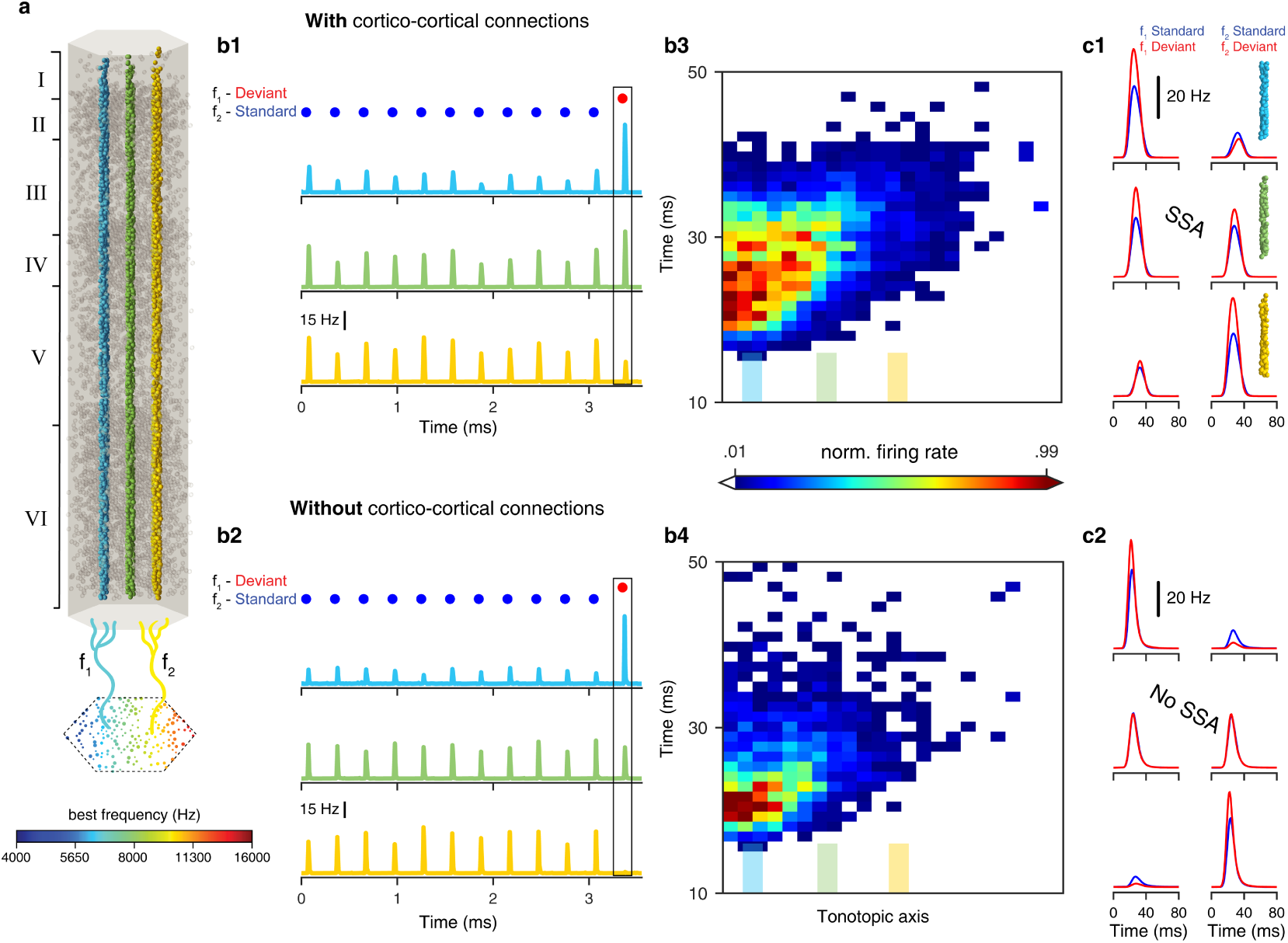
Combined effect of cortico-cortical connections and spike frequency adaptation on the emergence of SSA, in the absent of synaptic depression. **a.** Three reference “microcolumns” whose neurons respond best to different frequencies: Blue neurons receiving strong thalamic input for *f*_*1*_ = 6,666 Hz and weak thalamic input for *f*_*2*_ = 9,600 Hz; Green neurons receiving similar thalamic input for *f*_*1*_ and *f*_*2*_; Yellow neurons receiving strong thalamic input for *f*_*2*_ and weak thalamic input for *f*_*1*_. **b1**. Mean response of the cells in the three microcolumns shown in a when cortical connections remain untouched. **b2**. Without cortical connections, the strong response of the blue cells does not propagate to the green cells (compare green peak in the rectangle in **b2** vs. **b1**). In both cases, we removed synaptic depression from the circuit. **b3, b4**. Propagation of activity in the microcircuit for cases shown in the rectangles in **b1** and **b2**, respectively. The x-axis denotes left-to-right direction in the microcircuit shown in A; the locations of the 3 microcolumns are denoted by the filled colored bars above the x-axis. Note the extended spatial propagation right wise of the response for the case when *f*_*1*_ is the deviant for the connected circuit (compare **b4** to **b3**). Bin size – 1.2 ms over 15 µm. **c1, c2**. Left column: mean response of the three microcolumns shown in A for *f*_*1*_ as the standard (blue) and as deviant (red). The colored microcolumns at right mark the respective microcolumns shown in a. Right column: same as the left column but for *f*_*2*_. **c1** shows the case with cortico-cortical connections and **c2** the case without cortico-cortical connections.

We found that the neurons in the center microcolumn responded significantly more to the deviant tone than to the same tone when standard (**Fig. 2a1** and **a2**, red bars vs. blue bars). This behavior is the hallmark of the stimulus-specific adaptation^1^, showing that the model indeed succeeds to replicate the phenomenon. We validated that the center microcolumn is not selective to tone identity by showing that the cells respond equally to *f*_*1*_ and *f*_*2*_ when they are presented with equal probability (**Fig. 2a3**; the “equal paradigm”^14^, see additional quantification in **Supplementary Fig. 2**).

We next quantified the SSA in single excitatory cortical neurons; two examples are shown, one for Layer 2/3 pyramidal cell and one for Layer 5 pyramidal cell (**Fig. 2b1-b4**). In both neurons, the lower was the probability of the presented frequency, the higher was the response of the neuron (**Supplementary Fig. 2**). This behavior matched the results found *in vivo*^1,14,35^. To quantify the degree of SSA, we computed the common-contrast SSA index CSI^1,13^ (see **Methods**) for each neuron. CSI = 0 indicates that there is no difference in the response to the two frequencies when standard and deviant (**Fig. 2c**), while positive values indicate a stronger responses to at least one of the two tones when deviant. The CSI of most neurons was positive and so was the mean across layers (thick line). Note that the largest CSI was at middle layers and lower layer 6 which, interestingly, received relatively weak thalamic input (**Fig. 1b** and see **Discussion**). The average CSI value across all layers was 0.49 ± 0.29 (mean ± std). We also computed the SSA index (see **Methods**) for *f*_*1*_ (SI1) and for *f*_*2*_ (SI2) and plotted them in **Fig. 2d** against each other. Note that most values fall in the first quadrant, implying that the neurons are more responsive to the deviant than to the standard, irrespective of the tone frequency. The mean SSA index value was 0.47 ± 0.33 and 0.53 ± 0.39 for SI1 and SI2, respectively (red circle **Fig. 2d**). A summary of the SSA in the microcircuit for the present paradigm together with an additional stimulus paradigm is presented in **Supplementary Fig. 3** showing that, without any parameter tuning, the NMC model replicates rather successfully diverse aspects of SSA. Specifically, that the response of the neurons is significantly higher to the deviant than to the standard tone^1,3–5,13,14,29,31,35^, that this difference depends on the probability of the deviant^1,14,35^, and that the response to the deviant is larger than the response to the same tone when it is presented among many tones with densely-packed frequencies - the ‘diverse narrow’ paradigm^4,13,14^. Furthermore, in some cases, the response for a rarely presented tone (‘deviant alone’ or ‘rare’) were smaller than the responses to the same tone presented as a deviant accompanied by a standard tone^4,13,14^. However, **Supplementary Fig. 3d-f** shows that other experimental feature of SSA was not replicated by our model (see **Discussion**).

### Uncovering the neuronal mechanisms underlying SSA

#### The contribution of synaptic depression and spike-frequency adaptation to SSA

As our dense replica of a cortical microcircuit together with its biologically constrained set of mechanisms replicated many of the properties of SSA, we next used it to uncover the biophysical mechanisms underlying this phenomenon. Synaptic depression is widely considered as the fundamental mechanism underlying SSA. In the NMC it is modeled as a short-term reduction of the probability of a spike triggering synaptic transmission, as described in the Tsodyks-Markram model^36^. We therefore first removed this effect by fixing the release probability (both in the cortico-cortical and the thalamo-cortical synapses) of all excitatory depressing synapses (see **Methods**). Following this manipulation, the SSA was indeed reduced (compare **Fig. 3a** to **3b**). The mean CSI was reduced from 0.49 ± 0.29 (mean ± std) to 0.16 ± 0.2 (a 67% reduction) whereas the mean SI1 and SI2 indices changed from 0.47 ± 0.33 and 0.53 ± 0.39 to 0.16 ± 0.29 and 0.14 ± 0.33, respectively (a 69% reduction, compare red circle in insets in **Fig. 3a** to that of **Fig. 3b**). However, surprisingly, blocking synaptic depression did not completely eliminate SSA (**Fig. 3b**), implying that additional mechanisms support SSA in the model.

We therefore systematically examined the effect of a broad range of model parameters on the emergence of SSA. Including removing from the model a variety of membrane ion channels, simulating only some, out of the six, cortical layers, removing synaptic facilitation, etc. Eventually, we found that, when removing both synaptic depression and spike frequency adaptation (SFA), the SSA was completely abolished (**Fig. 3d**). SFA was removed by blocking Ca^2+^-dependent potassium channels from all modeled cells (see **Methods, Supplementary Fig. 4** and **Discussion**). Like synaptic depression, removing SFA alone reduced SSA but did not abolish it (**Fig. 3c**). Indeed, in this case, the mean CSI value was reduced from 0.49 ± 0.29 to 0.21 ± 0.29 (by 57% reduction). This results clearly shows that SSA emerges due to at least two mechanisms, synaptic depression and spike frequency adaptation, operating synergistically. The interaction between these two mechanisms is supralinear; while the average CSI value was 0.16 without synaptic depression (but with SFA) and 0.21 without SFA (but with synaptic depression), it was 0.49 when both mechanisms are present. For 1,719 out of 2,378 neurons, the summed CSI was smaller than the CSI for both mechanisms simultaneously (**Supplementary Fig. 5**).

The summary of the above results is depicted in **Fig. 3e** where the mean CSI value across cortical layers for the different conditions is shown. Note that, synaptic depression alone generates SSA (light green bars), and SFA alone generates SSA (orange bars). When the two mechanisms are active together, the SSA is larger (dark green bars). A detailed exploration of the origin of the differences in SSA among the different cortical layers is elaborated in the **Discussion**.

We next explored the contribution of cortico-cortical versus thalamo-cortical synaptic depression to SSA. We found that, in our model, thalamo-cortical synaptic depression has a much stronger impact on SSA than cortico-cortical synaptic depression. (Compare **Supplementary Fig. 6** first column, lower left frame to the frame in 2^nd^ row and 2^nd^ column). When only cortico-cortical depression is active (i.e., without SFA and thalamo-cortical synaptic depression) cortical neurons have a wide tuning curve. This is because, in this condition, the cortical neurons do not adapt (no SFA) and, on top of it, there is no thalamo-cortical synaptic depression^37^. Consequently, cortical neurons that are selective to, for example, *f*_*1*_ also respond strongly to *f*_*2*_. When *f*_*1*_ arrives as the deviant, their cortico-cortical synapses are already significantly depressed. In contrast, thalamic axons are narrowly tuned (responding selectively either to *f*_*1*_ or to *f*_*2*_). Consequently, in this condition, thalamic axons that are selective to *f*_*1*_ will respond weakly to *f*_*2*_. When *f*_*1*_ arrives as the deviant their thalamo-cortical synapses are only weakly depressed, resulting in a strong response to the deviant tone and, thus, with a strong SSA. Exploration of additional combinations of different mechanisms underlying SSA is elaborated in **Supplementary Fig. 6**.

### Emergence of SSA due to spike frequency adaptation

**Figure 3b** shows that, even in the absence of synaptic depression, SSA still persists in the microcircuit due to SFA (**Fig. 3d**). In the absence of synaptic depression, the cells consisting of the “center microcolumn”, i.e., cells that have no thalamo-cortical preference for either of the two stimuli (green cells in **Fig. 4a**), receive the same number of active thalamic inputs when *f*_*1*_ or *f*_*2*_ are presented. Therefore, on the face of it, these cells should respond in the same manner (and similarly adapt due to SFA) to *f*_*1*_ and *f*_*2*_. However, these “green neurons” also receive cortico-cortical inputs that affect their responses to *f*_*1*_ and *f*_*2*_. To test whether these cortico-cortical inputs mediate the effects of SFA on SSA, we removed all cortico-cortical synapses from the model. This manipulation, on top of removing synaptic depression (but leaving SFA intact), abolished SSA in these neurons (**Supplementary Fig. 6** first row in the 3^rd^ column). Thus, cortico-cortical connectivity was required in order for SFA to contribute to SSA.

To explore the role of cortico-cortical connectivity, we examined the responses of two additional microcolumns (blue and yellow cells in **Fig. 4a**) in the SSA protocol, in the absence of synaptic depression. The blue microcolumn consists of neurons that are selective for *f*_*1*_ (6,666 Hz); the yellow microcolumn is selective for *f*_*2*_ (9,600 Hz). The green microcolumn whose preferred frequency lies midway between *f*_*1*_ and *f*_*2*_ (8,000 Hz). When *f*_*2*_ was the standard, the *f*_*2*_-selective (“yellow” cells) adapted due to SFA (compare **Supplementary Fig. 3c**, “deviant alone” green bar, to the “standard”, blue bar). As a result, their feedforward effect on the green cells was small. When *f*_*1*_ was presented as deviant (in the same sequence in which *f*_*2*_ was standard), the *f*_*1*_-selective cells (“blue” cells) respond strongly (because they were not adapted by the many presentations of *f*_*2*_; **Fig. 4b1**, top). This effect on yellow and blue cells does not require cortico-cortical connectivity (**Fig. 4b1** vs. **4b2**, top), but in order for them to affect the activity of the green cells, cortico-cortical connections are required (**Fig. 4b1** vs. **4b2**, middle and **4b3** vs. **4b4**). Thus, in this condition, the green cells responded more strongly to the deviant (*f*_*1*_) than to the standard (*f*_*2*_). The reverse also holds for *f*_*2*_ as deviant and *f*_*1*_ as standard (**Fig. 4c1**, middle row, red trace is higher than the blue traces). This explains how SFA in conjunction with recurrent network connectivity causes SSA in the green cells. This cascade is illustrated with the animation in Supplementary Video 1. Remarkably, this situation recapitulates essential elements of the simplified model for SSA that is based on adaptation of narrowly tuned inputs^13,14^.

In conclusion, we note that, while, classically, synaptic depression was considered to be the major mechanism responsible for SSA, our dense *in silico* neocortical microcircuit model, receiving tonotopically-mapped thalamic input, enabled us to demonstrate that at least two additional mechanisms (SFA combined with network connectivity) are involved in shaping cortical SSA. Our study also shows that in our model SSA is primarily a network phenomenon because, when the network is disconnected, the thalamo-cortical input *per se* (and its respective synaptic depression) gives rise to only a small SSA response and only in neurons that receive thalamo-cortical inputs (**Supplementary Fig. 6** first row in 2^nd^ column).

### Additional properties of SSA explained via the reconstructed microcircuit

We next attempted to replicate, using the NMC model, and mechanistically explain other experimentally observed properties of SSA. One such property is the dependency of SSA on the amplitude (intensity) of the presented tones. It was found experimentally that stronger tones result in lower CSI^29,30^. We replicated this experiment by setting the firing rate of the modeled thalamic axons so that it mimics the experimental response to different tone intensities (Sound Pressure Levels, SPL, **Fig. 5a, Supplementary Fig. 1**, and **Methods**). Indeed, as found experimentally, the microcircuit showed a decrease in CSI with the increase in tone intensity (**Fig. 5b**). When examining the frequency-response area (FRA) of thalamic axons (**Fig. 5a**), we observe that two main features of the tuning curve change with tone intensity. The higher the tone intensity, the higher is the firing rate of the thalamic axons and the wider is their tuning curve (**Fig. 5a** and **Supplementary Fig. 1**). To separate the effect of each of these two parameters on CSI we manipulated, in the model, either the tuning width of the thalamic axons (**Fig. 5c**) or their maximal firing rate (**Fig. 5d**), leaving all other variables unchanged (see **Methods**). The range of values used for testing the effects of the width of the tuning curve (**Fig. 5c**) and of the firing rate (**Fig. 5d**) was within the experimental range (**Supplementary Fig. 1**). Both the increase in firing rate and in tuning width resulted in a decrease in the CSI, but the change in the tuning width had a much greater effect (compare **Fig. 5c** to **5d**).

**Figure 5.**
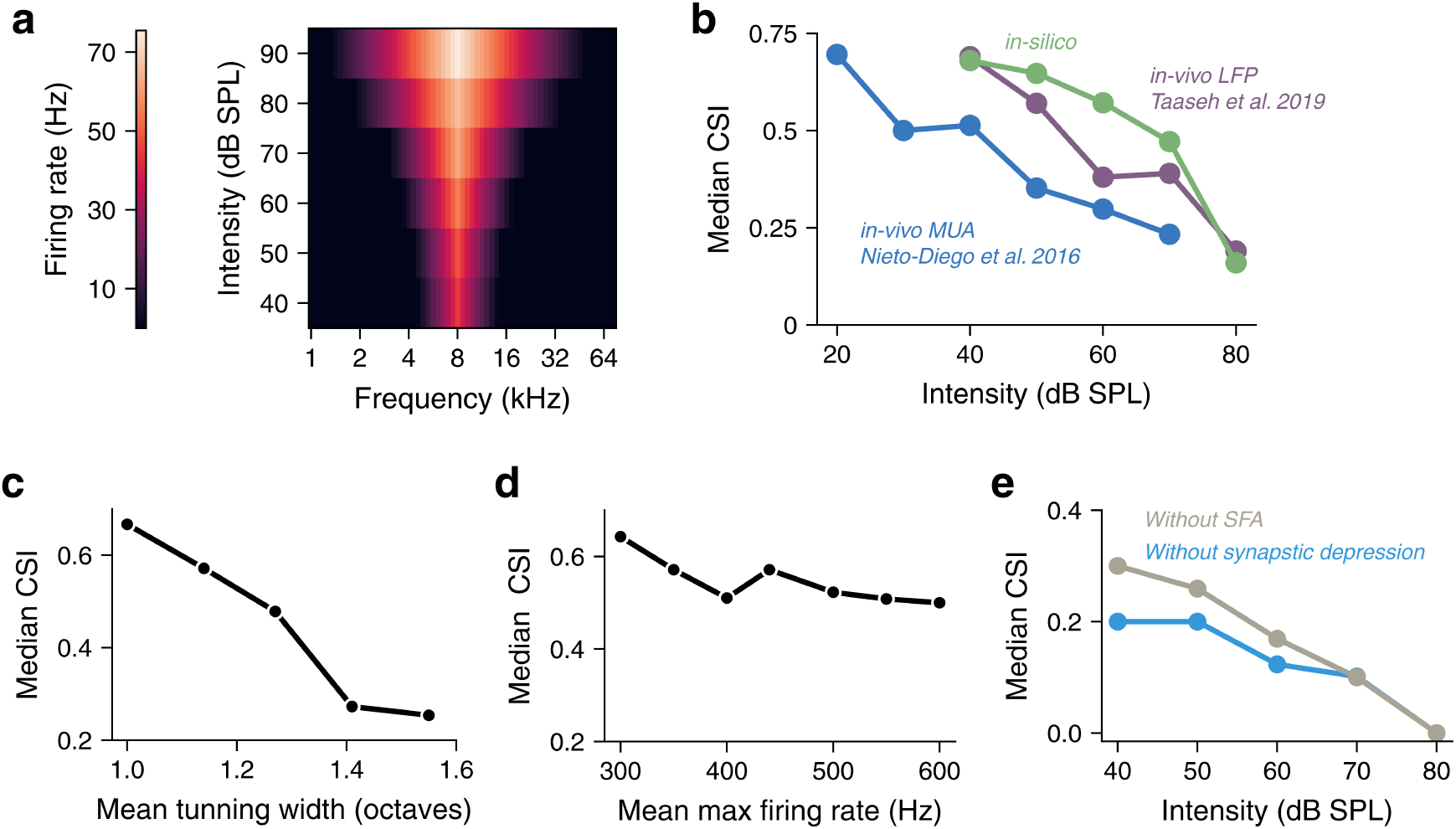
Effect of tone intensity on SSA. **a**. Mean, experimentally based, frequency response area (FRA) for the modeled thalamic axons when the preferred frequency of each axon is shifted to 8 kHz (see **Supplementary Fig. 1** and **Methods**). **b**. Median CSI value as a function of tone intensity measured from the 2,696 modeled neurons as in **Fig. 2a** (green line), superimposed on two *in vivo* experimental results (blue and purple lines). **c, d**. Effect of changing the width of the tuning curve (**c**) and the maximal firing rate (**d**) on the CSI while keeping all other parameters fixed as in the response to 60 dB (see **Methods** and **Supplementary Fig. 1**). **e**. Effect of removing spike frequency adaptation (grey line) or synaptic depression (blue line) on the dependency of CSI on tone intensity.

Next, we utilized the circuit to uncover which of the two biophysical mechanisms (synaptic depression or SFA) is responsible for the reduction in CSI with increasing tone intensity. To achieve this, we replicated the protocol of **Fig. 5b**, once when removing SFA and once when blocking synaptic depression (**Fig. 5e**). Both manipulations resulted in a comparable reduction of the CSI with an increase in tone intensity (**Fig. 5e**), implying that both mechanisms are involved in the dependence of CSI on tone intensity. This is to be expected, because when tone intensity is increased, the thalamo-cortical afferents selective to the deviant also start responding to the standard due to the increase in tuning width. As a consequence, both synaptic adaptation and SFA will reduce the strength of the response to the deviant as a consequence of exposure to the standard. Consequently, in cases with high tone-intensity, the difference in response between a standard and a deviant is smaller (smaller CSI).

Finally, we explored the experimental finding that the strength of the SSA depends on the frequency preference of the cortical neurons^31^. Indeed, the tuning curve of neurons in the auditory cortex is typically not symmetrical and, therefore, the neurons do not respond with the same firing rate to the two tones (*f*_*1*_ and *f*_*2*_) that are centered around their best frequency^10,13,29,31^. Chen et al. (2015) reported that, when using the oddball paradigm, the SI of the tone that gave rise to the lower firing rate (the “Non-Preferred” frequency SI_NP_; e.g., *f*_*2*_,) was lower than the SI of the tone that gave rise to the higher firing rate (the “Preferred” frequency SI_P_; e.g., *f*_*1*_; See **Fig. 5** in^31^). We independently verified this result by analyzing the dataset accompanying Nieto-Diego et al., 2016 (see **Methods**). We found that SI_P_ is significantly larger than SI_NP_ (0.221 ± 0.035 versus 0.11 ± 0.037, n = 31; p = 0.032 t-test; mean ± SEM). **Fig. 6a, b** shows that this also occurs in our simulated microcircuit. We first plotted the firing rate of the excitatory cortical neurons as a function of the tonotopic axis (**Fig. 6a**). As expected, cortical neurons that are closer to the thalamic axons with *f*_*1*_ as the BF (black circle) have a higher firing rate for *f*_*1*_ (black line in **Fig. 6a**) and *vice versa* for *f*_*2*_ (grey line in **Fig. 6a**). **Fig. 6b** shows that, as in the experiment, the larger the firing rate of a neuron to *f*_*2*_ relative to *f*_*1*_ the larger the SI2 is (and vice versa for SI1, not shown). Together, **Fig. 6a** and **6b** correspond to the experimental finding that the tone that gives rise to the lower firing rate has lower SI value.

**Figure 6.**
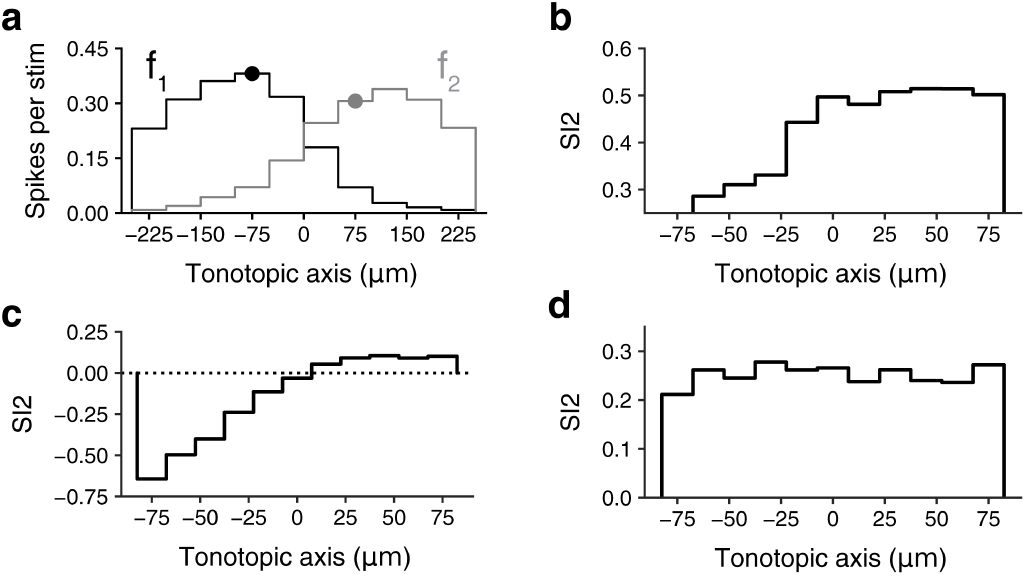
Dependence of SSA on the frequency preference of the cell. **a**. Mean number of spikes of the excitatory neurons in the simulated microcircuit in response to *f*_*1*_ (6,666 Hz, black line) or *f*_*2*_ (9,600 Hz, grey line) when played as the standard, as a function of the tonotopic axis (left-to-right direction in the microcircuit shown in **Fig. 4a**; bin size: 50 µm). The black and grey circles mark the position of the thalamic axons with BF of *f*_*1*_ and *f*_*2*_. **b**. SI2 as a function of the tonotopic axis. Cortical neurons that are closer to the thalamic axons with BF of *f*_*2*_ (grey circle in a) have higher SI2 (bin size: 15 µm). **c**. SI2 in a model circuit leaving only spike frequency adaptation (cortico-cortical connections and thalamic synaptic depression are removed). The less responsive the neuron is to *f*_*2*_ (progressing left along the tonotopic axis, see a) the lower the SI2 is. **d**. SI2 in the model circuit when only SFA was removed, demonstrating that, in this case, the dependency of SI2 on the tonotopic axis (on tone frequency) vanishes.

This finding can be understood by considering the involvement of SFA in the emergence of SSA (**Fig. 3d**). In the absence of cortico-cortical connectivity, SFA may cause a neuron that is selective to *f*_*1*_ to respond more strongly to *f*_*2*_ as standard than to *f*_*2*_ as deviant (low SI2). This may happen when *f*_*1*_ is highly effective in driving the neuron, and therefore, when standard, the firing of this neuron would be strongly adapted. Consequently, when *f*_*2*_ occurs as a deviant, the response of the neuron is small. If *f*_*2*_ is much less effective in driving the neuron, SFA would affect the neuronal responses to a lesser extent and the neuron would respond more strongly to *f*_*2*_ under this condition. Thus, for this cell, SI2 is expected to be low and possibly even negative. We tested that prediction by removing cortico-cortical connections and synaptic depression from the model and found that indeed neurons that are more selective to *f*_*1*_ have lower SI2 (**Fig. 6c**). Furthermore, we found that, when blocking SFA, while leaving all other mechanisms intact, the dependence of SI2 on the frequency preference of the neuron vanishes (**Fig. 6d**).

## Discussion

This work proposes a general approach, which could be termed “replicate and explain”, for linking experimentally-observed (emergent) circuit responses to their underlying anatomical and biophysical properties. This approach utilizes a generic dense computer replica of the circuit studied as an “experimental” tool that incorporates a broad range of biophysical and anatomical details (membrane ion channels, synaptic and firing dynamics, cell and synapse types, dendritic morphology, cortical layers, etc.). Yet, this circuit model was not built to capture any *a priori* specific high-level phenomenon. We have used the “replicate and explain” approach to study SSA in the auditory system, utilizing the cortical microcircuit model of young rat developed by Markram et al., (2015), and adding to this circuit tonotopically mapped thalamo-cortical axons^33,34^ together with modeling the response of these axons to pure tones based on our own unpublished experimental data (**Supplementary Fig. 1**). This dense circuit replicated a substantial amount (but not all, see below) of the experimental properties of SSA, without tuning of the underlying modeling parameters; it provided mechanistic explanations for the emergence of SSA and related phenomena, together with a set of experimentally-testable predictions. The study suggested that a significant part of SSA is not a special property of the whole brain *in vivo*, nor is it specific to the auditory cortex and it has provided a set of new mechanisms that support SSA.

Our “replicate and explain” approach differs from the typical computational route taken to study circuit dynamics, whereby simplified circuit models are constructed to study the dependence of an observed phenomenon, e.g., SSA, on unobserved biophysical parameters^38,39^, for example, synaptic depression. While this classical modeling approach is invaluable for testing specific mechanisms that might explain the observed phenomena, the model is *a priori* tailored to replicate the phenomenon and, by design, it ignores other putative underlying mechanisms (and the interactions among them) that might impact the studied phenomena^40^.

In agreement with previous theoretical suggestions, we found that thalamo-cortical synaptic depression (and to a lesser degree also cortico-cortical synaptic depression) plays a key role in generating SSA^13,15^. Yet, removing synaptic depression from the modeled circuit uncovered additional mechanisms responsible for SSA. Namely, a synergetic “collaborative” interaction between spike frequency adaptation, SFA, at the cellular level and cortico-cortical network connections. SFA was previously proposed as a putative mechanism underlying SSA^41^, but its impact at the network level for generating SSA was not demonstrated. We have shown that SFA, in the absence of network connectivity, does not generate SSA in neurons that are equally selective to *f*_*1*_ and *f*_*2*_ (**Fig. 4**), whereas in neurons that are not equally selective to *f*_*1*_ and *f*_*2*_, SFA results with positive SI for the frequency (say *f*_*1*_) that the neuron is selective for and negative or zero for *f*_*2*_ (**Fig. 6**). This is in contrast to the experimental observation, whereby both SI1 and SI2 were positive^13^. One key experimentally testable prediction from this study is that SSA will be significantly weakened by blocking ion channels responsible for SFA (i.e., the Ca^2+^-dependent K channels, which could be blocked by albumin^42^). This study thus shows how a computation (in our case of a “surprise response”) can be generated in networks receiving spatially mapped sensory input (e.g., tonotopic input, retinotopic input) via SFA mechanism.

In accordance with experimental results^3,43^ we found that SSA was weaker in the thalamo-recipient layers (lower layers 3 and 5 in our model, see **Fig. 1b** and **2c**). In the thalamo-recipient layers the standard tone generated a strong response and a weaker response in the other layers. When the deviant arrives, neurons in all layers increase their firing rate by approximately the same amount. This implies that the CSI (and thus SSA) will be lower in the thalamo-cortical recipient layers (see **Methods** for CSI definition). Note also that neurons in the non-recipient layers (that respond weakly to a standard tone) are less adapted and this allows them to increase their response more significantly to the deviant tone. Our “replicate and explain” methodology explained two additional SSA-related experimental phenomena. First, by modeling *in vivo* spike time recordings of MGBv neurons responding to pure tones at various sound pressure levels (SPL), we tested the oddball paradigm at different SPL. We found that, as in the experiments, the larger the tone intensity the lower the SSA^29,30^. The main reason for this effect is the widening of the thalamic axon tuning curve with tone intensity which, in turn, affects the capability of both SFA and synaptic depression to generate SSA. Second, we found that, in agreement with the experiments, SSA depended on the frequency preference of the neurons^31^ and showed that SFA is the main mechanism responsible for this dependency (**Fig. 6**).

Our model did not succeed in replicating the experimental distribution of SSA values in different cortical layers (compare **Fig. 2c** to **Fig, 7** in ^3^) nor did it replicated an important property of SSA in primary auditory cortex: its specific deviance sensitivity. In auditory cortex, responses to a tone when deviant (accompanied by a standard) are comparable in size and sometimes larger than the responses to the same tone, occurring at the same probability, when accompanied by many widely-spaced, tones. This is the ‘control’ condition of the mismatch negativity in EEG literature^8^ and the ‘diverse broad’ condition in^13,14^. In our model, the responses to a deviant frequency were almost always smaller than the response to this frequency in the ‘diverse broad’ condition (see **Supplementary Fig. 3d-f**). Thus, the deviant responses, while larger than standard responses, are still too small to fit the experimentally measured ones. This is actually also the major failure of previous models that were based on adaptation of narrowly-tuned inputs^6,13,14^. This might be the result of our ability to model only six tones in the diverse broad condition, compared to the twelve tones that are used experimentally. We also note that this response depends on the location of the neurons, where neurons with BS close to the deviant frequency show a deviant response that is larger than the response to the ‘diverse broad’ (**Supplementary Fig. 3e**), although only for one frequency. Further modeling efforts are required to solve this puzzling experimental result.

With the increase in computer power we are able, for the first time, to connect via a dense computer replica of neuronal circuits, the subcellular (ion channels and synapses), the cellular (cell types) and the network (connectivity) levels, and obtain a new and lucid picture of the working of the brain at multitude levels of operation^44^. This will enable to resolve other long-standing open questions such as the emergence of “salt and pepper” organization in sensory systems of rodents^45–48^ and the impact of different excitatory and/or inhibitory cell types on network dynamics^5,49–51^. Furthermore, this approach could be scaled-up for asking as broader questions as we will soon have detailed computer replicas of the rodent hippocampus, cerebellum, thalamus, as well as a detailed model of the whole rodent neocortex with full thalamo-cortical connections^52–55^. Then we might explore, and provide mechanistic explanations, for a wide range of *in vivo* multiscale phenomena such as EEG signals (and its modulation, e.g., due to a “surprise”) which, after more than 100 years since they were discovered, are only partially understood

## Methods

### Dense model of neocortical microcircuit (NMC)

We performed simulations of electrical activity on a previously published model of a neocortical microcircuit of a two-week-old rat. Reconstruction of the circuits and its simulation methods were described in Markram et al.^19^. Briefly, this microcircuit (**Fig. 1a**) consisted of 31,346 biophysical Hodgkin-Huxley NEURON models and around 7.8 million synaptic connections forming around 36.4 million synapses. Synaptic connectivity between 55 distinct morphological types of neurons (m-types) was predicted algorithmically and constrained by experimental data ^24^. The densities of ion-channels on morphologically-detailed neuron models were optimized to reproduce the behavior of different electrical neuron types (e-types) and synaptic dynamics recorded *in vitro*^56^. Simulations were run on HPE SGI 8600 supercomputer (BlueBrain V) using NEURON^57^ and CoreNEURON^58^. NEURON models and the connectome are available online at bbp.epfl.ch/nmc-portal^59^. In **Supplementary Fig. 3d-f** a larger circuit was simulated (see below).

### Simulation of baseline spontaneous activity

To account for the missing long-range connections and missing neuromodulators, neurons were depolarized with a somatic current injection of 85% of first spike threshold. In addition, synapses spontaneous release probability was set to mimic an *in vivo* like state. As described by Markram et al.^19^, the use parameter of synaptic transmission and the spontaneous release probability were modulated according to extracellular calcium concentration ([Ca^2+^]_o_). In all our simulations [Ca^2+^]_o_ was set to 1.23mM (see also ^28^). Synaptic conductances and kinetics are as in Markram et al.^19^. For the excitatory, synapses, the NMDA component was revised such that the “steepness” factor, γ, of the NMDA conductance was set to 0.08^60–62^.

### Simulation of realistic auditory input delivered to the NMC

Auditory input was simulated using 574 reconstructed thalamic axons^19^ that were activated tonotopically as follows: Each thalamic axon had x, y coordinates which correspond to the center of its projection to the NMC. The x-coordinate spans from 0 to 560 µm and was denoted as the tonotopic axis (**Fig. 1d**). The best frequency, BF, of each axon corresponds linearly to its x-coordinate, such that BF of 4,000 Hz corresponded to x = 0 µm, and BF of 16,000 Hz to x = 560 µm. This process resulted in 2 octaves difference along the x-axis^33,34,63,64^.

To set the firing-rate dynamics of the thalamic axons so that it replicates the firing-rate dynamics of the rodent auditory thalamic nucleus, we first analyzed *in vivo* recordings of rat ventral division of the medial geniculate body (MGBv), (see below). We examined the response of MGBv neurons to pure tones at frequencies ranging from 1 kHz to 64 kHz (steps of 0.166 octaves) and at amplitudes ranging from 40 to 90 dB Sound Pressure Levels (SPL). Out of 26 MGBv neurons recorded experimentally, 17 had a V-shaped frequency response area (FRA) when measured in the first 50 ms after tone presentation (see example in **Supplementary Fig. 1a** left). We then used the instantaneous firing rate of each of these 17 MGBv neurons as the target for the fit using Eq. (1), which is a standard alpha function (see example in **Supplementary Fig. 1b** and see also ^65^). The other 9 of the axons were either non-responsive to tone presentation or had a highly irregular tuning curve. Their mean firing-rate was 1.5 Hz.

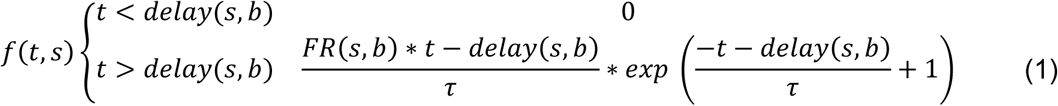

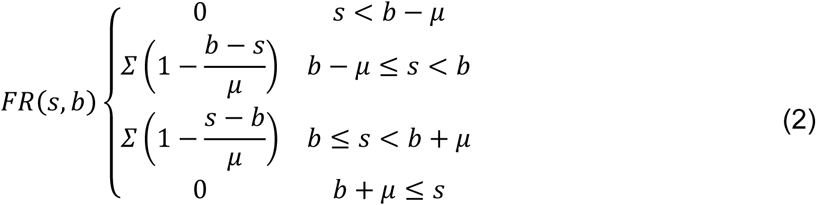

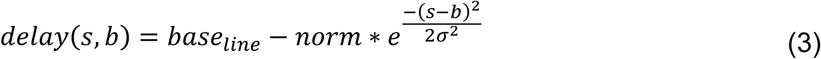

Where *s* is the presented frequency and *t* is time. The parameters for this alpha function are: *b* - the neuron best frequency; ∑ - maximum firing rate; *µ* - width of tuning curve; *τ* - response time constant. The frequency-dependent delay (delay(*s,b*)) had a Gaussian waveform (e.g., the delay is shortest when *s* = *b)* with the following parameters: *base*_*line*_ - the maximal delay; *norm -* the max delay difference; *σ* - the Gaussian standard deviation. These parameters were fitted with SciPy^66^ least-square function (using the default parameters) with the target of minimizing ∑_*s*_ f(t, s) − MGBv(t, s). This process resulted in a distribution for each of parameter, for each tone amplitude (**Supplementary Fig. 1c-h**). We used these parameter distributions to create the spike times of the thalamic axons.

Because 9 out of 26 MGBv axons did not have a characteristic V-shape response, we started by randomly selecting 40% of the modeled thalamic axons and activated them spontaneously at the rate to 1.5 Hz (see above). For each of the remaining (60%) thalamic axons we randomly sampled the parameters ∑ and *τ* from a Gaussian distribution that fitted the corresponding parameter distribution in **Supplementary Fig. 1c-h** and *µ* from a 2-parameter exponential distributed that fitted its distribution (**Supplementary Fig. 1c-h**). The *base*_*line*_, *norm, σ* that were selected for the delay function were that of the mean of the corresponding parametric distribution (**Supplementary Fig. 1c-h**). Using this sampling process, we satisfy all parameters for *f*(*t, s*). Then, for each tone presentation, the spike times of the axon were generated from an inhomogeneous Poisson process with time-varying rate *f*(*t, s*) + 1.5 Hz.

In **Figure 5c**,**d** all the parameters for each axon were sampled from the parameter distribution for an input intensity of 60 dB (**Supplementary Fig. 1e**) except for the location parameter of the exponential distribution for the tuning width (**Supplementary Fig. 5c**) and for the mean of the Gaussian distribution for the max firing rate (**Supplementary Fig. 5d**).

The FRA in **Figure 5a** was constructed by creating a thalamic axon with the average values of ∑, *τ* and *µ* for each tone amplitude, and averaging the first 50 ms of its response to different tone presentations.

### SSA protocols

We attempted to replicate as closely as possible the *in vivo* experimental protocols used to explore SSA in rodent auditory cortex^4,5,13,14,29,35,67^. To achieve this, two tones *f*_*1*_ = 6,666 Hz and *f*_*2*_ = 9,600 Hz were selected as the base frequencies for our simulations; these frequencies are separated by 0.52 octaves around the center frequency of the simulated cortical circuit (8,000 Hz); see **Fig. 1d** and **Fig. 4a**. Each simulated experiment started with 2 seconds without an auditory tone followed by 500 tone presentations, with an inter-stimulus interval (ISI) of 300 ms. We presented tones in five paradigms, described here. The oddball paradigm, where the standard tone was presented at p = 95% (unless specified otherwise) and the deviant tone was presented with p = 5%. The order of the tone presentations was random. In the equal paradigm, each tone was presented 250 times in a random order. In the deviant alone paradigm, the deviant timing was identical to the deviant timing in oddball paradigm, but the standard tone was not presented at all. In the diverse narrow paradigm, 20 tones equally distributed between *log*_2_(*f*_1_) − 0.19 *to log*_2_(*f*_2_) + 0.19 were each presented 25 times in a random order. In the diverse broad paradigm – 6 tones equally distributed between *log*_2_(*f*_1_) − 1.05 *to log*_2_(*f*_2_) + 1.05. For simulating this condition (**Supplementary Fig. 3d**), we simulated a 3D-slice of cortical circuit containing ∼103,207 neurons consisting of 1000 µm along the tonotopic (x-axis) by 465 µm width (y-axis) and with same depth (z-axis) as in as in **Fig. 1a**. This 3D-slice was taken from the full 7-column circuit in Markram et al.^19^.

### Data analysis

The analysis of SSA response in the circuit was conducted on neurons at ± 25 µm about the center of the simulated circuit. These neurons have a best frequency of around 8,000 Hz and respond similarly to *f*_*1*_ and *f*_*2*_ when each is presented alone (see **Supplementary Fig. 3b1**). Unless stated otherwise, common-contrast SSA Index (CSI) measurements (and SI1, SI2; see below) are shown only for neurons with a firing rate larger than 0.15 Hz. For the oddball paradigm, we compared the neuron response (50 ms from tone presentation) for 24 randomly selected standards (*f1s* or *f2s*) to their response for 24 deviants (*f1d* or *f2d*). We discarded the first response to a standard and a deviant tone

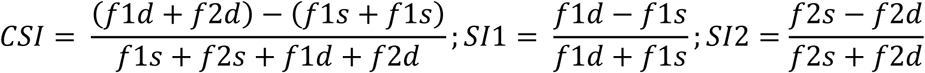

### Manipulation of synaptic depression

Synapses in the microcircuit are separated into three types of short-term synaptic plasticity: facilitating, depressing, and pseudo-linear. The dynamics of these synapses is governed by the respective set of time constants^19^. To remove short time synaptic plasticity from a given synaptic connection, we set the facilitation and depression time constants to 0 while keeping the release probability constant.

### Manipulation of spike frequency adaptation

Excitatory neuron in the microcircuit contains calcium-activated potassium channels (SK_E2, see also ^68,69^). This hyperpolarizing current was shown to be involved in generating spike frequency adaptation, SFA^70^. To remove the SFA from the microcircuit we abolished this current by removing all SK_E2 channels from all modeled compartments.

### Analysis of previously published data

The experimental plots in **Supplementary Fig. 4c** (black and gray lines) were digitally reconstructed from **Fig. 3d** in Abolafia et al., 2011, and overlaid on the axes of the simulated results.

The experimental data from Taaseh, 2010, in **Fig. 5b** (purple line) was digitally extracted (5 median points) from **Fig. 5d** of Yarden and Nelken^11^

The experimental data from Nieto-Diego et al.^29^, in **Fig. 5b** (blue line) was created by analyzing the dataset accompanying the article, S1 Data Oddball (MUA). We extracted all SSA results of A1 units and calculated the median CSI as a function of SPL.

We used the same dataset to calculate SI_P_ and SI_NP_. For each neuron that was recorded in SPL>50, we compared its firing rate response to *f*_*1*_ and to *f*_*2*_ as standards. The SI of the frequency that gave rise to the larger response was defined as SI_P_ whereas the SI of the frequency that gave rise to the smaller response was defined as SI_NP._

### Visualization

Network visualizations were created in **Fig. 1b** using RTNeuron^71^ and in **Fig. 1c** using Mayavi^72^. Single neuron morphology visualizations (**Fig. 2b**) were created using NeuroMorphoVis^73^. Figures were created using Matplotlib^74^.

### Experimental procedures

The thalamic inputs were modeled based on unpublished data recorded in rat MGB, courtesy of Leila Khouri and Maciej Jankowski. All experiments were performed according to protocols approved by the Institutional Animal Care and Use Committee of Hebrew University. Hebrew University is an AAALAC approved institution. Animal preparation was as in Taaseh et al., 2011. Following the craniotomy, an electrode array (16 or 32 tungsten electrodes, Alpha-Omega) was introduced to the MGB using stereotaxic coordinates. The data analyzed here consisted of responses to pure tones of 37 different frequencies (1,000 Hz to 32,000 Hz, 6 tones/octave, 50 ms long with 10 ms linear onset and offset ramps, 2/s). Each tone was repeated 10 times at each sound level. Sound levels covered the range between 20-30 dB SPL to 90 dB SPL at 10 dB steps. All data were stored as disk files and analyzed offline (Matlab, Mathworks). Following the experiment, the rats were perfused transcardially, the brains removed and used for histological verification of the recording locations. Units with large spikes were isolated using a unit-specific threshold. Only units with clear significant responses to tones were used.

## Acknowledgments

We thank Pramod Kumbhar for the dedicated technical support. We thank all members of the Segev labs for many fruitful discussions and valuable feedback regarding this work. We thank Leila Khouri and Maciej Jankowski for the experimental data.

## Funding

This study received funding from the European Union’s Horizon 2020 Framework Program for Research and Innovation under the Specific Grant Agreement No. 785907 (Human Brain Project SGA2), the ETH domain for the Blue Brain Project, the Gatsby Charitable Foundation, the NIH Grant Agreement U01MH114812 and support by a personal ISF grant (1126/18).

## Competing financial interests

Authors declare no competing financial interests.

## Data and materials availability

All data will be published upon publication

## Authors Contribution

O.A. and I.S. conceived the study. O.A., I.S, I.N, M.R and H.M wrote the manuscript. O.A. carried out the simulations and the analysis. I.N contributed the experimental data. H.M., S.R., J.K., M.R, E.M, O.A. and I.S developed the *in silico* microcircuit and provided the respective simulation data.

## Supplementary Information

**Supplementary Figure 1.**
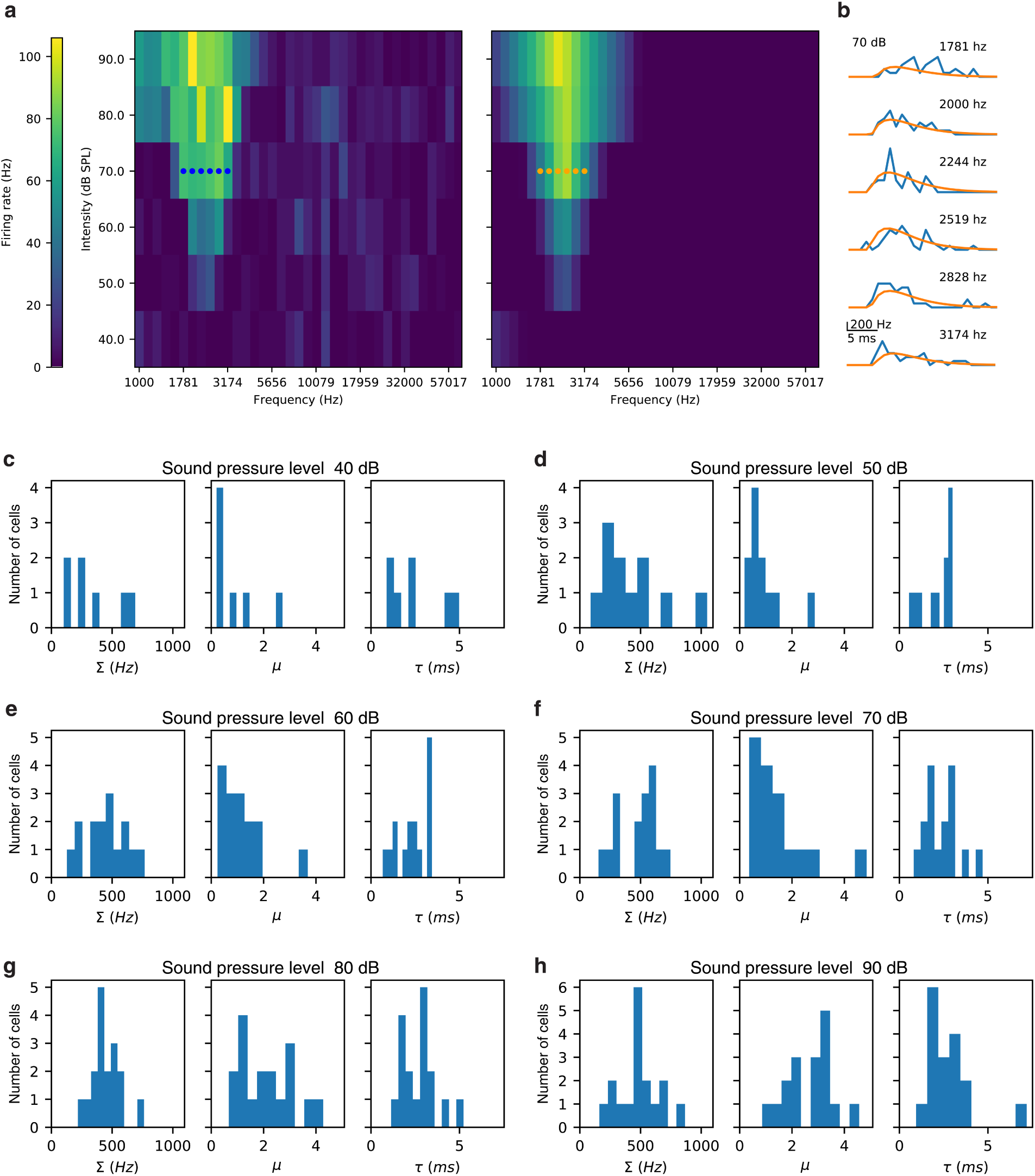
Fitting experimental responses of MGBv neurons for generating auditory thalamic input. **a**. Exemplary FRA from an MGBv thalamic neuron (left) and the FRA for the fit to the experimental data (right). Bin size – 0.167 octaves measured on the first 50 ms after tone presentation. **b**. The experimental and fitted instantaneous firing rate for 6 different frequencies for an amplitude of 70 dB (marked by blue and orange points in a. **c-h**. Distributions of the values that were fitted for the different MGBv neurons (variables fit of Eqs (1-2); see **Methods**)

**Supplementary Figure 2.**
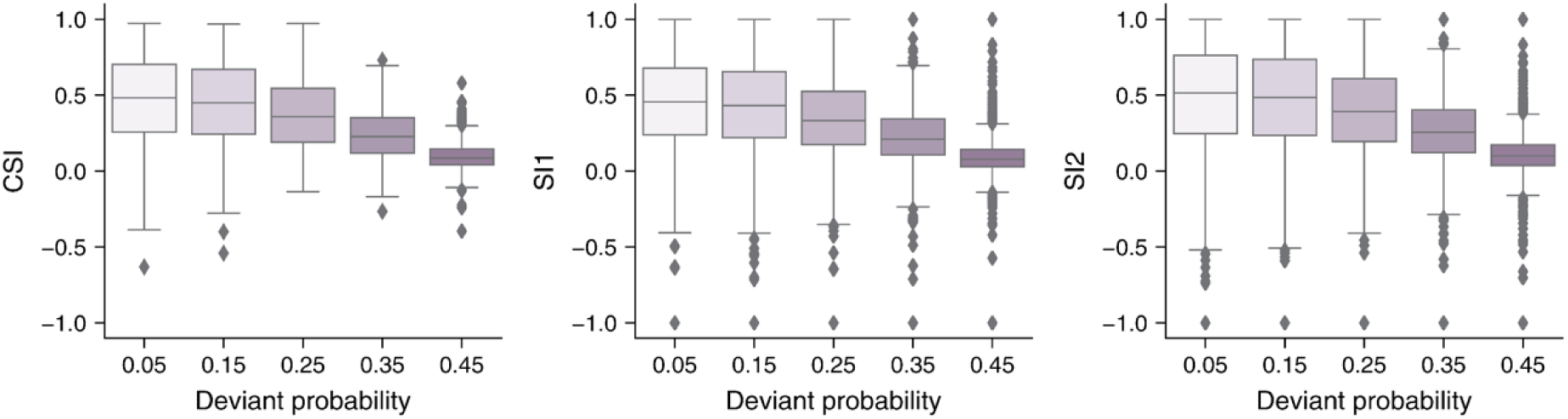
SSA dependence of the probability of the presented tone. The lower the probability of the deviant tone in the oddball paradigm, the higher is the SSA (quantified by CSI, SI1, and SI2, see **Methods**).

**Supplementary Figure 3.**
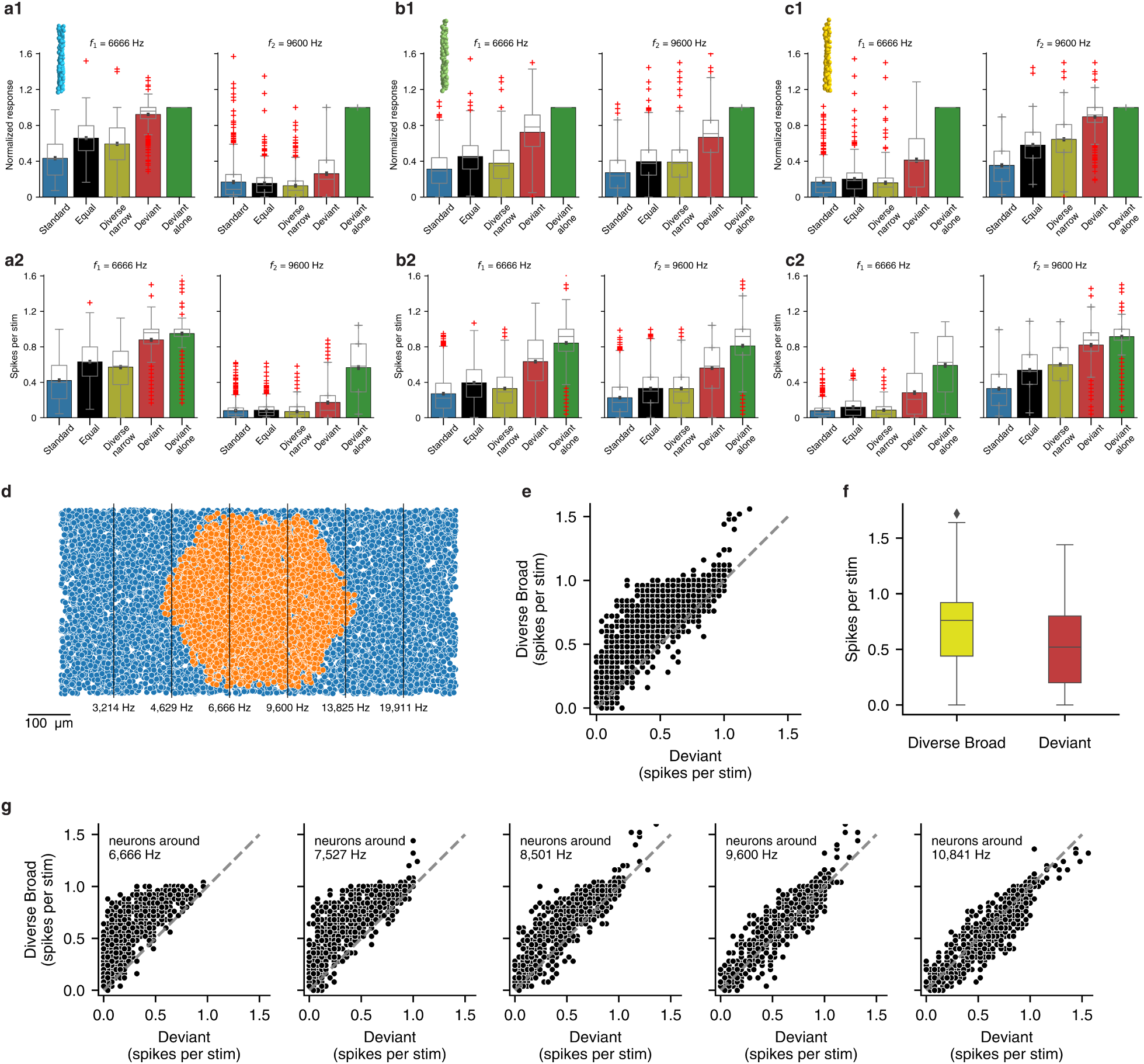
Neurons response to additional auditory SSA protocols. **a1-c1**. Response amplitude (spikes per tone presentation) for multiple experimental paradigms for the blue cells (**a1**) green cells (**b1**) and yellow cells (**c3**) in **Fig. 4a. a2-c2.** Same as a1-c1 but when normalized by the corresponding response for the deviant alone paradigm. **d.** Microcircuit used for the diverse broad protocols (microcircuit used for all other figures in this study is marked in orange). Black lines mark the location of the 6 frequencies that were used. **e.** Spikes per tone when *f*_*1*_ is presented as deviant and when presented in a diverse broad protocol (response of 2,000 cells are shown by points). **f.** Box plot for the data shown in e. **g.** Same as e but for neurons with different locations (±25 μm around the BF marked in the legend)

**Supplementary Figure 4.**
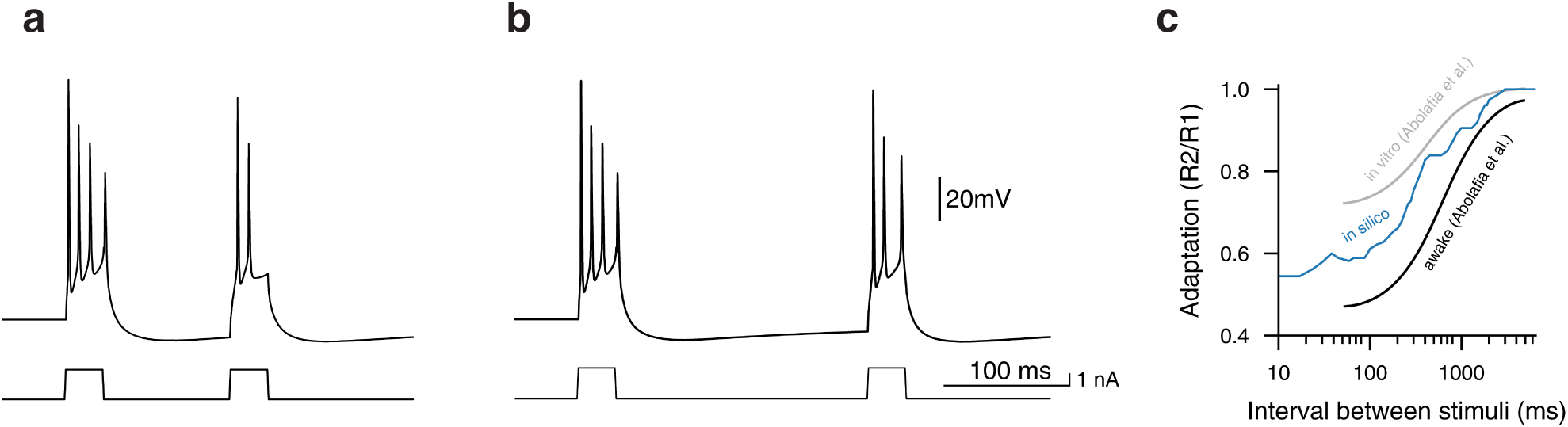
Spike frequency adaptation. **a**. Example of adaptation protocol for a layer 5 TTPC2 extracted from the microcircuit. Current injection was set to five times rheobase current (2.5 nA) and ISI to 100 ms. Note the decrease in the number of spikes for the second current injection. **b**. Same as a but for a longer ISI (200 ms). **c**. Adaptation index, which is defined by the ratio between the number of spikes for the second current injection (R2) to the number of spikes for the first current injection (R1), as a function of the time between the current injections, for 25 excitatory cells (5 from each layer) from the microcircuit (blue line, *in silico*). Experimental results for *in vitro* and awake preparations are shown by the gray and black lines, respectively^1^.

**Supplementary Figure 5.**
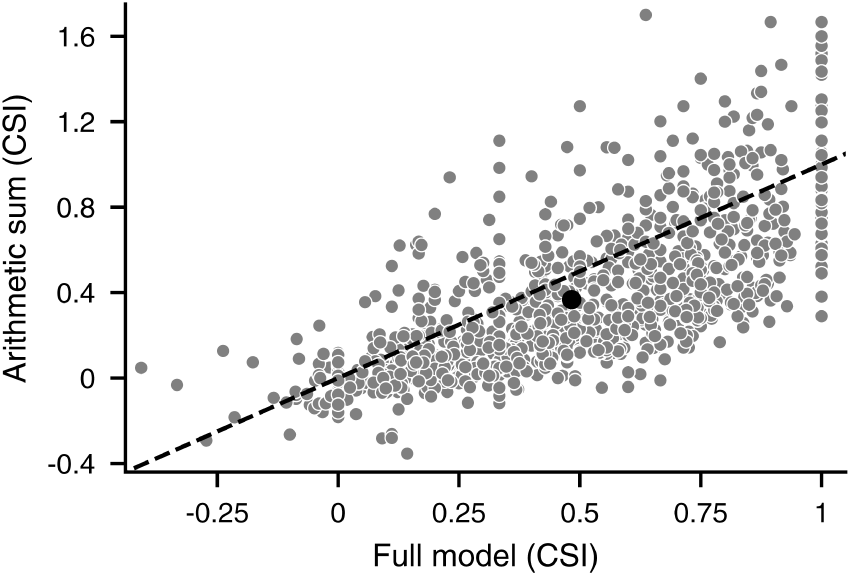
SSA supralinearity. CSI of 1000 excitatory cells measured from the full microcircuit (with both synaptic depression and SFA; **Fig. 3a**) versus the arithmetic sum of the respective CSI in the absent of synaptic depression (**Fig. 3b**) plus the CSI in the absent of SFA (**Fig. 3c**). Black dot marks the mean CSI.

**Supplementary Figure 6.**
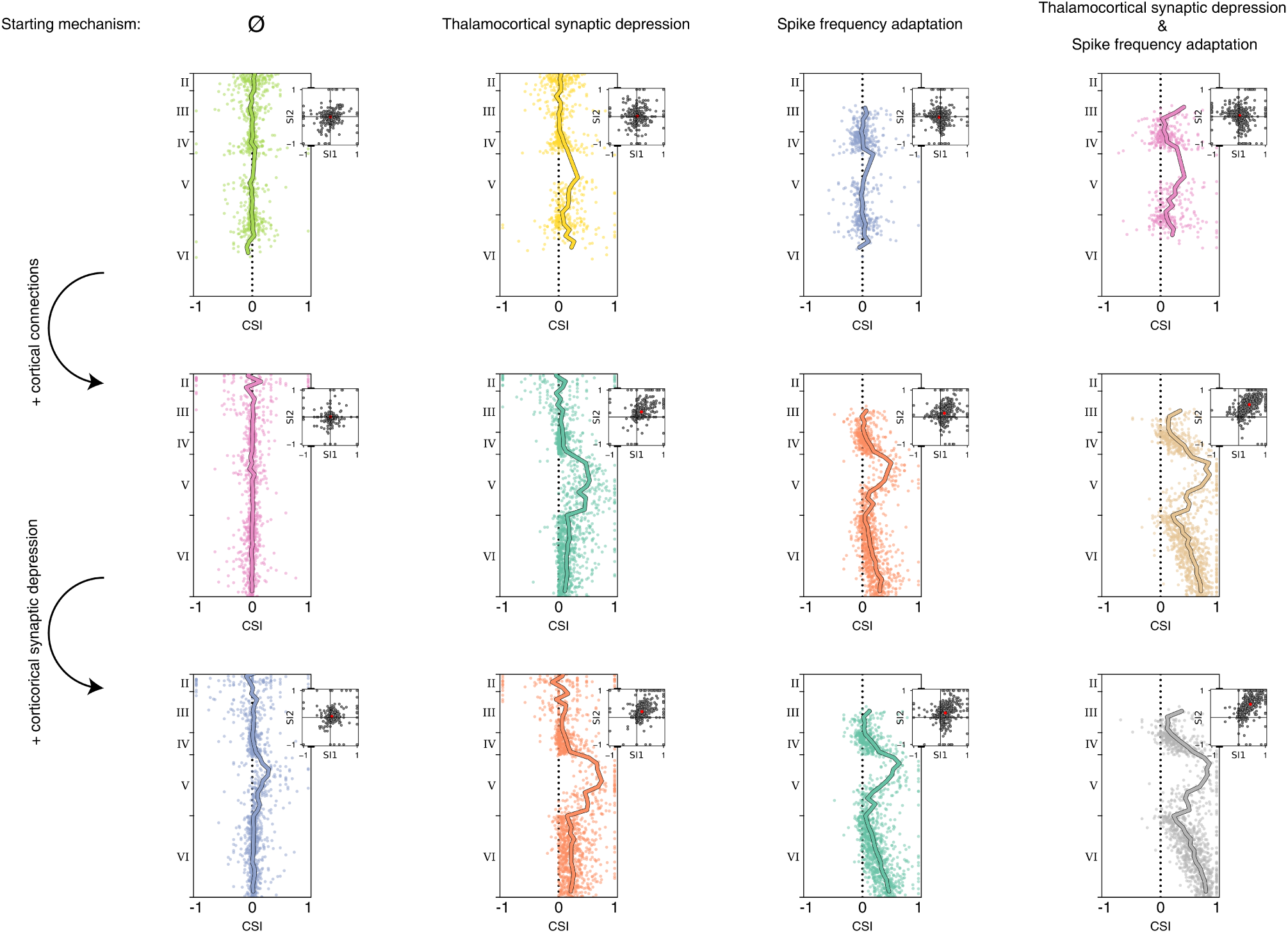
SSA strength over a variety of network configurations. Each panel in the first row presets the CSI values in a circuit with the mechanism/s titling the corresponding panel. The panels in the 2^nd^ row show the CSI values when cortical connections were added to the corresponding circuits in the first row. The panels in the 3^rd^ row show the CSI when cortical synaptic depression is added to the corresponding circuit in the 2^nd^ row.

## Notes

### Competing Interest Statement

The authors have declared no competing interest.

## References

1. Ulanovsky, N., Las, L. & Nelken, I. Processing of low-probability sounds by cortical neurons. Nat. Neurosci. 6, 391–8 (2003).

2. Von Der Behrens, W., Bäuerle, P., Kössl, M. & Gaese, B. H. Correlating stimulus-specific adaptation of cortical neurons and local field potentials in the awake rat. J. Neurosci. (2009) doi:10.1523/JNEUROSCI.3475-09.2009.

3. Szymanski, F. D., Garcia-Lazaro, J. A. & Schnupp, J. W. H. Current source density profiles of stimulus-specific adaptation in rat auditory cortex. J. Neurophysiol. 102, 1483–1490 (2009).

4. Polterovich, A., Jankowski, M. M. & Nelken, I. Deviance sensitivity in the auditory cortex of freely moving rats. PLoS One (2018) doi:10.1371/journal.pone.0197678.

5. Natan, R. G. et al. Complementary control of sensory adaptation by two types of cortical interneurons. Elife 4, e09868 (2015).

6. Nelken, I. Stimulus-specific adaptation and deviance detection in the auditory system: experiments and models. Biol. Cybern. 108, 655–63 (2014).

7. Nelken, I., Yaron, A., Polterovich, A. & Hershenhoren, I. Stimulus-Specific Adaptation Beyond Pure Tones. in 411–418 (Springer, New York, NY, 2013). doi:10.1007/978-1-4614-1590-9_45.

8. Jacobsen, T., Schröger, E., Horenkamp, T. & Winkler, I. Mismatch negativity to pitch change: Varied stimulus proportions in controlling effects of neural refractoriness on human auditory event-related brain potentials. Neuroscience Letters (2003) doi:10.1016/S0304-3940(03)00408-7.

9. Antunes, F. M., Nelken, I., Covey, E. & Malmierca, M. S. Stimulus-specific adaptation in the auditory thalamus of the anesthetized rat. PLoS One 5, e14071 (2010).

10. Ulanovsky, N., Las, L., Farkas, D. & Nelken, I. Multiple Time Scales of Adaptation in Auditory Cortex Neurons. J. Neurosci. 24, 10440–10453 (2004).

11. Yarden, T. S. & Nelken, I. Stimulus-specific adaptation in a recurrent network model of primary auditory cortex. PLoS Comput. Biol. 13, (2017).

12. Farley, B. J., Quirk, M. C., Doherty, J. J. & Christian, E. P. Stimulus-specific adaptation in auditory cortex is an NMDA-independent process distinct from the sensory novelty encoded by the mismatch negativity. J Neurosci 30, 16475–16484 (2010).

13. Taaseh, N., Yaron, A. & Nelken, I. Stimulus-specific adaptation and deviance detection in the rat auditory cortex. PLoS One 6, e23369 (2011).

14. Hershenhoren, I., Taaseh, N., Antunes, F. M. & Nelken, I. Intracellular Correlates of Stimulus-Specific Adaptation. J. Neurosci. 34, 3303–3319 (2014).

15. Mill, R., Coath, M., Wennekers, T. & Denham, S. L. A Neurocomputational Model of Stimulus-Specific Adaptation to Oddball and Markov Sequences. PLoS Comput. Biol. 7, e1002117 (2011).

16. Mill, R., Coath, M., Wennekers, T. & Denham, S. L. Characterising stimulus-specific adaptation using a multi-layer field model. in Brain Research vol. 1434 178–188 (2012).

17. May, P. J. C., Westö, J. & Tiitinen, H. Computational modelling suggests that temporal integration results from synaptic adaptation in auditory cortex. Eur. J. Neurosci. 41, 615–630 (2015).

18. Wacongne, C., Changeux, J.-P. & Dehaene, S. A Neuronal Model of Predictive Coding Accounting for the Mismatch Negativity. J. Neurosci. 32, 3665–3678 (2012).

19. Markram, H. et al. Reconstruction and Simulation of Neocortical Microcircuitry. Cell 163, 456–492 (2015).

20. Reyes-Puerta, V., Sun, J. J., Kim, S., Kilb, W. & Luhmann, H. J. Laminar and Columnar Structure of Sensory-Evoked Multineuronal Spike Sequences in Adult Rat Barrel Cortex in Vivo. Cereb. Cortex (2015) doi:10.1093/cercor/bhu007.

21. Renart, A. et al. The asynchronous state in cortical circuits. Science (80-.). (2010) doi:10.1126/science.1179850.

22. Luczak, A., Barthó, P., Marguet, S. L., Buzsáki, G. & Harris, K. D. Sequential structure of neocortical spontaneous activity in vivo. Proc. Natl. Acad. Sci. U. S. A. (2007) doi:10.1073/pnas.0605643104.

23. Okun, M. et al. Diverse coupling of neurons to populations in sensory cortex. Nature (2015) doi:10.1038/nature14273.

24. Reimann, M. W., Muller, E. B., Ramaswamy, S. & Markram, H. An Algorithm to Predict the Connectome of Neural Microcircuits. Front. Neural Circuits 9, (2015).

25. Reimann, M. W., Horlemann, A. L., Ramaswamy, S., Muller, E. B. & Markram, H. Morphological diversity strongly constrains synaptic connectivity and plasticity. Cereb. Cortex (2017) doi:10.1093/cercor/bhx150.

26. Gal, E. et al. Rich cell-type-specific network topology in neocortical microcircuitry. Nat. Neurosci. 20, 1004–1013 (2017).

27. Reimann, M. W. et al. Cliques of neurons bound into cavities provide a missing link between structure and function. Front. Comput. Neurosci. (2017) doi:10.3389/fncom.2017.00048.

28. Nolte, M., Reimann, M. W., King, J. G., Markram, H. & Muller, E. B. Cortical reliability amid noise and chaos. Nat. Commun. (2019) doi:10.1038/s41467-019-11633-8.

29. Nieto-Diego, J. et al. Topographic Distribution of Stimulus-Specific Adaptation across Auditory Cortical Fields in the Anesthetized Rat. PLoS Biol. 14, e1002397 (2016).

30. Taaseh, N. The Cellular Basis of Pre-Attentive Evoked Potentials. (The Hebrew University of Jerusalem, 2010).

31. Chen, I.-W., Helmchen, F. & Lütcke, H. Specific Early and Late Oddball-Evoked Responses in Excitatory and Inhibitory Neurons of Mouse Auditory Cortex. J. Neurosci. 35, 12560–73 (2015).

32. Meyer, H. S. et al. Cell Type–Specific Thalamic Innervation in a Column of Rat Vibrissal Cortex. Cereb. Cortex 20, 2287–2303 (2010).

33. Polley, D. B., Read, H. L., Storace, D. A. & Merzenich, M. M. Multiparametric Auditory Receptive Field Organization Across Five Cortical Fields in the Albino Rat. J. Neurophysiol. (2007) doi:10.1152/jn.01298.2006.

34. Kenet, T., Froemke, R. C., Schreiner, C. E., Pessah, I. N. & Merzenich, M. M. Perinatal exposure to a noncoplanar polychlorinated biphenyl alters tonotopy, receptive fields, and plasticity in rat primary auditory cortex. Proc. Natl. Acad. Sci. (2007) doi:10.1073/pnas.0701944104.

35. Yaron, A., Hershenhoren, I. & Nelken, I. Sensitivity to complex statistical regularities in rat auditory cortex. Neuron 76, 603–15 (2012).

36. Tsodyks, M. V & Markram, H. The neural code between neocortical pyramidal neurons depends on neurotransmitter release probability. Proc. Natl. Acad. Sci. U. S. A. 94, 719–23 (1997).

37. Banitt, Y., Martin, K. A. C. & Segev, I. A biologically realistic model of contrast invariant orientation tuning by thalamocortical synaptic depression. J. Neurosci. (2007) doi:10.1523/JNEUROSCI.1640-07.2007.

38. Sterratt, D., Graham, B., Gillies, A. & Willshaw, D. *Principles of Computational Modelling in Neuroscience*. Principles of Computational Modelling in Neuroscience (Cambridge University Press, 2011). doi:10.1017/CBO9780511975899.

39. Amsalem, O., Van Geit, W., Muller, E., Markram, H. & Segev, I. From Neuron Biophysics to Orientation Selectivity in Electrically Coupled Networks of Neocortical L2/3 Large Basket Cells. Cereb. Cortex 26, 3655–3668 (2016).

40. Fan, X. & Markram, H. A brief history of simulation neuroscience. Frontiers in Neuroinformatics (2019) doi:10.3389/fninf.2019.00032.

41. Abolafia, J. M., Vergara, R., Arnold, M. M., Reig, R. & Sanchez-Vives, M. V. Cortical auditory adaptation in the awake rat and the role of potassium currents. Cereb. Cortex 21, 977–990 (2011).

42. Pennefather, P., Lancaster, B., Adams, P. R. & Nicoll, R. A. Two distinct Ca-dependent K currents in bullfrog sympathetic ganglion cells. Proc. Natl. Acad. Sci. U. S. A. (1985) doi:10.1073/pnas.82.9.3040.

43. Musall, S., Haiss, F., Weber, B. & von der Behrens, W. Deviant Processing in the Primary Somatosensory Cortex. Cereb. Cortex 27, 863–876 (2017).

44. Einevoll, G. T. et al. The Scientific Case for Brain Simulations. Neuron (2019) doi:10.1016/j.neuron.2019.03.027.

45. Rothschild, G., Nelken, I. & Mizrahi, A. Functional organization and population dynamics in the mouse primary auditory cortex. Nat. Neurosci. 13, 353–360 (2010).

46. Ohki, K., Chung, S., Ch’ng, Y. H., Kara, P. & Reid, R. C. Functional imaging with cellular resolution reveals precise microarchitecture in visual cortex. Nature (2005) doi:10.1038/nature03274.

47. Fahey, P. G. et al. A global map of orientation tuning in mouse visual cortex. bioRxiv (2019) doi:10.1101/745323.

48. Maor, I., Shalev, A. & Mizrahi, A. Distinct Spatiotemporal Response Properties of Excitatory Versus Inhibitory Neurons in the Mouse Auditory Cortex. Cereb. Cortex (2016) doi:10.1093/cercor/bhw266.

49. Lee, S.-H., Kwan, A. C. & Dan, Y. Interneuron subtypes and orientation tuning. Nature 508, E1–2 (2014).

50. Cardin, J. A. et al. Driving fast-spiking cells induces gamma rhythm and controls sensory responses. Nature 459, 663–7 (2009).

51. Adesnik, H. & Scanziani, M. Lateral competition for cortical space by layer-specific horizontal circuits. Nature 464, 1155–1160 (2010).

52. Markram, H. et al. Reconstruction and simulation of a scaffold model of the cerebellar network. Front. Neuroinform. 163, 456–92 (2019).

53. Reimann, M. W. et al. A null model of the mouse whole-neocortex micro-connectome. Nat. Commun. (2019) doi:10.1038/s41467-019-11630-x.

54. Iavarone, E., Yi, J., Shi, Y., Markram, H. & Hill, S. L. Digital reconstruction and simulation of thalamic microcircuitry. in Society for Neuroscience Program No. 340.08 (2019).

55. Billeh, Y. N. et al. Systematic Integration of Structural and Functional Data into Multi-Scale Models of Mouse Primary Visual Cortex. SSRN Electron. J. (2019) doi:10.2139/ssrn.3416643.

56. Van Geit, W. et al. BluePyOpt: Leveraging Open Source Software and Cloud Infrastructure to Optimise Model Parameters in Neuroscience. Front. Neuroinform. 10, 17 (2016).

57. Carnevale, N. T. & Hines, M. L. The NEURON Book. (Cambridge University Press, 2006). doi:10.1017/CBO9780511541612.

58. Kumbhar, P. et al. CoreNEURON : An Optimized Compute Engine for the NEURON Simulator. Front. Neuroinform. (2019) doi:10.3389/fninf.2019.00063.

59. Ramaswamy, S. et al. The neocortical microcircuit collaboration portal: a resource for rat somatosensory cortex. Front. Neural Circuits 9, 44 (2015).

60. Poleg-Polsky, A. Effects of neural morphology and input distribution on synaptic processing by global and focal NMDA-spikes. PLoS One (2015) doi:10.1371/journal.pone.0140254.

61. Rhodes, P. The properties and implications of NMDA spikes in neocortical pyramidal cells. J. Neurosci. (2006) doi:10.1523/JNEUROSCI.3791-05.2006.

62. Doron, M., Chindemi, G., Muller, E., Markram, H. & Segev, I. Timed Synaptic Inhibition Shapes NMDA Spikes, Influencing Local Dendritic Processing and Global I/O Properties of Cortical Neurons. Cell Rep. 21, 1550–1561 (2017).

63. Zhou, X. & Merzenich, M. M. Enduring effects of early structured noise exposure on temporal modulation in the primary auditory cortex. Proc. Natl. Acad. Sci. U. S. A. (2008) doi:10.1073/pnas.0800009105.

64. Tischbirek, C. H. et al. In Vivo Functional Mapping of a Cortical Column at Single-Neuron Resolution. Cell Rep. 27, 1319-1326.e5 (2019).

65. Larsson, J. P. Modelling neuronal mechanisms of the processing of tones and phonemes in the higher auditory system. (Universitat Pompeu Fabra, 2013).

66. Virtanen, P. et al. SciPy 1.0--Fundamental Algorithms for Scientific Computing in Python. (2019).

67. Parto Dezfouli, M., Zarei, M., Jahed, M. & Daliri, M. R. Stimulus-specific adaptation decreases the coupling of spikes to LFP phase. Front. Neural Circuits 13, 1–11 (2019).

68. Kohler, M. et al. Small-Conductance, Calcium-Activated Potassium Channels from Mammalian Brain. Science (80-.). 273, 1709–1714 (1996).

69. Hay, E., Hill, S., Schürmann, F., Markram, H. & Segev, I. Models of neocortical layer 5b pyramidal cells capturing a wide range of dendritic and perisomatic active properties. PLoS Comput. Biol. 7, e1002107 (2011).

70. Engel, J., Schultens, H. A. & Schild, D. Small Conductance Potassium Channels Cause an Activity-Dependent Spike Frequency Adaptation and Make the Transfer Function of Neurons Logarithmic. Biophys. J. 76, 1310–1319 (1999).

71. Hernando, J. B., Biddiscombe, J., Bohara, B., Eilemann, S. & Schürmann, F. Practical Parallel Rendering of Detailed Neuron Simulations. in Eurographics Symposium on Parallel Graphics and Visualization (eds. Marton, F. & Moreland, K.) (The Eurographics Association, 2013). doi:10.2312/EGPGV/EGPGV13/049-056.

72. Ramachandran, P. & Varoquaux, G. Mayavi: 3D visualization of scientific data. Comput. Sci. Eng. (2011) doi:10.1109/MCSE.2011.35.

73. Abdellah, M. et al. NeuroMorphoVis: A collaborative framework for analysis and visualization of neuronal morphology skeletons reconstructed from microscopy stacks. in Bioinformatics (2018). doi:10.1093/bioinformatics/bty231.

74. Hunter, J. D. Matplotlib: A 2D Graphics Environment. Comput. Sci. Eng. 9, 90–95 (2007).

